# Small GTPases and BAR domain proteins regulate branched actin to make clathrin and dynamin independent endocytic vesicles

**DOI:** 10.1101/170753

**Authors:** Mugdha Sathe, Gayatri Muthukrishnan, James Rae, Andrea Disanza, Mukund Thattai, Giorgio Scita, Robert G. Parton, Satyajit Mayor

## Abstract

Numerous endocytic pathways operate simultaneously at the cell surface. Here we focus on the molecular machinery involved in the generation of endocytic vesicles of the clathrin and dynamin-independent CLIC/GEEC (CG) pathway. This pathway internalises many GPI-anchored proteins and a large fraction of the fluid-phase in different cell types. We developed a real-time TIRF assay using pH-sensitive GFP-GPI to identify nascent CG endocytic sites. The temporal profile of known CG pathway modulators showed that ARF1/GBF1 (GTPase/GEF pair) and CDC42 (RhoGTPase) are recruited sequentially to CG endocytic sites, ∼60s and ∼9s prior to scission. Using a limited RNAi screen, we found several BAR domain proteins affecting CG endocytosis and focused on IRSp53 and PICK1 that have interactions with CDC42 and ARF1 respectively. IRSp53, an I-BAR domain containing protein, was recruited to the plasma membrane at the site of forming CG endocytic vesicles and in its absence, nascent endocytic CLICs, did not form. The requirement for actin polymerization in the CG pathway suggested a role for nucleators of actin polymerization, and ARP2/3 was found enriched at the site of the forming endocytic vesicle. PICK1, a BAR domain containing protein and the ARP2/3 inhibitor is recruited at an early stage along with ARP2/3, but is removed from the endocytic site coincident with CDC42 recruitment and a burst of Factin polymerization. This study provides a spatio-temporal understanding of the molecular machinery necessary to build a CG endocytic vesicle.

## Introduction

Multiple endocytic pathways function in a eukaryotic cell^1,2^, however, our understanding of the endocytic process is mainly derived from studies on **C**lathrin-**M**ediated **E**ndocytosis (CME)^3–6^. Dynamin is responsible for vesicle scission in CME^7,8^ and a host of **C**lathrin-**I**ndependent **E**ndocytic (CIE) pathways, such as the caveolar and fast endophilin-mediated endocytic pathway^9–11^. On the other hand, among CIE pathways, the <u>**C**</u>LIC/<u>**G**</u>EEC [CLathrin and dynamin-Independent Carriers which form GPI-Enriched Endocytic Compartments; CG] pathway functions independent of both clathrin and dynamin in multiple cell types and contexts^12–18^, while it is not present in others^19^. The actin-polymerization machinery has been implicated in the functioning of many CIE pathways at different stages^14,20^.

Our focus, the CG pathway, which is regulated by the small GTPases, ARF1 (ADPribosylation factor 1) and CDC42 (Cell division control protein 42)^12–17^ is responsible for the uptake of many GPI-anchored proteins, a major fraction of the fluid-phase, toxins such as *Helicobacter pylori* vacuolating toxin A^21^, cholera toxin^22^ and viruses like Adeno-associated virus 2^23^. The CLICs are formed in a polarised manner at the leading edge of migrating cells^18^ and, the resulting GEECs subsequently fuse with the sorting endocytic vesicles via a Rab5/phosphatidylinositol-3-kinase dependent mechanism^24^. The CLICs/GEECs are high capacity endocytic carriers turning over the entire membrane surface in 12 minutes in fibroblasts, highlighting the role of CG pathway in regulating membrane homoeostasis^18^. Recent evidence suggests that this is required for generating a tubular vesicular endocytic network during cytokinesis^25^ and serves to deliver ligands to their signalling receptors in a common endocytic compartment.

The molecular machinery to form a CG endocytic vesicle involves activating ARF1 at the plasma membrane by GBF1 (Golgi-specific brefeldin A resistance factor 1)^17^, a specific ARF-GEF (Guanine nucleotide exchange factor). GTP-ARF1 recruits ARHGAP21 (a RhoGAP for CDC42) which removes CDC42 from the membrane^15^. Cholesterol removal, in addition, reduces the recruitment of ARF1 and CDC42, along with accelerated cycling of CDC42^14,15^. Lastly, the CG pathway requires dynamic actin since both stabilisation and depolymerization of actin filaments was found to affect endocytosis^14^.

By visualising a forming CG endocytic vesicle we wanted to understand the molecular mechanism responsible. We adapted a pH-pulsing protocol that exploits the pH-sensitive properties of Super Ecliptic GFP^26^, previously deployed to study CME^5,27,28^. We tagged the GPI anchor with Super Ecliptic GFP-GPI (SecGFP-GPI) to assay, in real-time, the sites of endocytic vesicle formation. We found that the CG endocytic vesicle formation is initiated by the accumulation of ARF1/GBF1 followed by CDC42 and F-actin while dynamin and clathrin did not associate with forming endosomes. Hence, in the absence of a discernable coat^18^, Bin/Amphiphysin/Rvs (BAR) domain containing proteins (BDPs) by generating/stabilising membrane curvature^29,30^ can assist in endocytic vesicle formation.

Several BDPs are involved in the CME pathway whereas only one has been identified to be associated with the CG endocytic pathway^31^. Using RNAi-screening, we identified 2 BDPs in particular, that affected CG endocytosis downstream of ARF1 and CDC42. Firstly, a CDC42 interaction partner and I-BAR protein, IRSp53 (Insulin-responsive protein of mass 53 kDa) was found to be necessary for CG endocytosis. Importantly, IRSp53 removal resulted in the disappearance of CLICs and loss of a GBF1-dependent endocytic pathway. Secondly, an ARF1 interaction partner, PICK1 (protein interacting with C kinase 1) emerged as an ARP2/3 activity regulator in the early phases of CG vesicle formation. Lastly, ARP2/3, an interaction partner of both IRSp53 and PICK1, accumulated at the forming CG endocytic site and decreased CG endocytosis when inhibited. Together, the spatio-temporal dynamics of these proteins provided a mechanistic understanding of the forming CG endocytic vesicle.

## RESULTS

### Identification of newly formed endocytic vesicles of the CG pathway containing GPI-anchored proteins

To monitor endocytic vesicle formation in real-time we employed the pH-sensitive fluorescence of Super Ecliptic pHlourinGFP^26^ attached to a GPI-anchor (SecGFP-GPI) to differentiate cell surface resident molecules from the newly internalised molecules. The fluorescence of SecGFP fluorescence is quenched reversibly when exposed to pH 5^5,26–28^. SecGFP-GPI expressed in AGS (human gastric cell line) was endocytosed along with the CG cargo, fluid-phase (10 kDa dextran), but not with CME cargo, TfR, at both 37°C and 30°C (Figure S1a-c & Supplementary Information (S.I.)) shown previously^16,32^. Endocytic events were identified by alternately exposing the cells to buffers equilibrated to pH 7 or pH 5 every 3s at 30°C (Figure 1a, Schematic & Supplementary Movies 1-2). SecGFP-GPI-containing endocytic events occurring during exposure to pH 7 remained fluorescent due to their neutral luminal pH right after formation. However, the buffer exchange from pH 7 to pH 5 quenched the fluorescence of cell surface SecGFP-GPI. This enabled visualisation of the newly formed endocytic vesicle. The identification of the site of endocytic vesicle formation paved the way for the characterization of the spatial and temporal dynamics of molecular players by the co-expression of an (mCherry/TagRFPt/pRuby)-tagged molecule of interest, ‘X-FP’ (Figure 1a & Supplementary Movies 1-2). The dynamics of X-FP was extracted by looking at the history of the region where vesicle formation was detected (Figure 1a montage (bottom), See S.I.).

**Figure 1:**
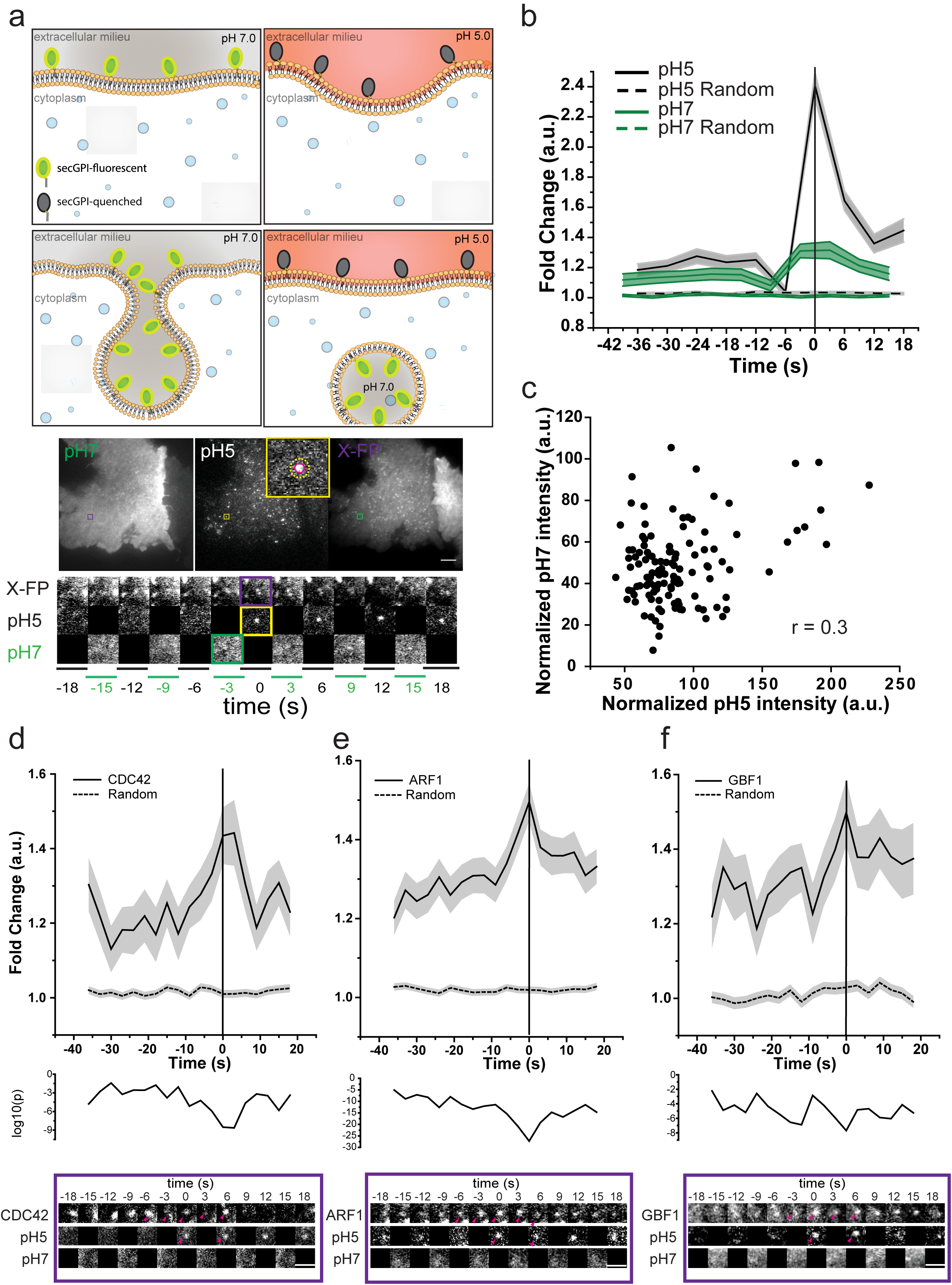
Identification of newly formed SecGFP-GPI endocytic vesicles using a pH pulsing assay. (**a**) Schematic (top panel) of the pH pulsing assay depicting the effect of pH on SecGFP-GPI fluorescence during endocytic vesicle budding. Note the fluorescence of SecGFP-GPI is retained at high pH (top and bottom left panel) or when exposed to low pH if sequestered in a newly formed endocytic vesicle (bottom right panel), and quenched only when exposed to low pH (top right panel). Sequential TIRF images of AGS cell co-expressing SecGFP-GPI and mCherry-ARF1 collected at pH 7, pH 5 and in the RFP channels (middle panel). Newly formed endocytic vesicles (inset) (identified as in Supplementary **Figure S1c**), are used construct a single frame (yellow rectangle) of the montage depicted (bottom panel). (**b**) Average of the normalized fluorescence intensities of pH 5 and pH 7 traces at the site of newly formed SecGFP-GPI endocytic vesicles compared to their respective random traces constructed from 120 endocytic events (pH5, pH7) and 3428 random spots, derived from 17 cells pooled from 4 independent experiments. (**c**) Graph shows the fold enrichment of fluorescence intensity over local background of pH 5 vs. pH 7 at the time of formation of the endocytic vesicles (data from 1b). (**d-f**) Graphs show the average normalized fluorescence intensity versus time traces for the recruitment of TagRFPtCDC42 (**d**), mCherry-ARF1 (**e**) and mCherry-GBF1 (**f**) to the forming SecGFP-GPI endocytic sites, and its corresponding random intensity trace (n, Table 2). The random traces were derived from randomly assigned spots of the same radius as the endocytic regions, as detailed in S.I. Endocytic distribution at each time point was compared to the random distribution by Mann-Whitney U test, and the log_10_ (p) [log_10_ (0.05) is −1.3 and log_10_ (0.001) is −2.5] is plotted below each trace (**d-f**). Representative montages from the respective data sets are depicted below the graphs (**d-f**). Arrowheads indicates the newly formed endocytic vesicle. Error bars, s.e.m (**b**,**d-f**). Scale bar, 1.5µm (**a**,**d-f**).

The pH pulsing movies were analysed using semi-automated scripts (See S.I.). Briefly, the centroid of new spots appearing in the pH 5 channel provided a fiduciary marker for the time and location of the nascent endocytic vesicle (Figure S1d, Step 1 & S.I.). The relative enrichment of SecGFP-GPI and X-FP at the endocytic site was determined by normalising the average fluorescence of the nascent endocytic spot to its local background annulus (Figure 1a, S1d, Step 2-3 & See S.I.). The spots were then put through a series of automated and manual checks. The automated check ensured, that the pH 5 intensity of the new spot had i) significantly higher intensity than the background, ii) persisted for at least 6s, and iii) did not show an increase in intensity in the subsequent frame. Subsequently, a manual check was performed on the montages, i) to remove any false positives that might have been missed by the automated check and ii) to classify the new SecGFP-GPI spots into two groups based on whether X-FP co-detection was observed or not (See S.I.). The data at the site of the spot was represented as the average fold change over the surrounding background, as a function of time (Figure 1b, solid traces), and compared to the average fold change of arbitrary regions within the cell (‘Random’) (Figure 1b, dashed traces). The profile obtained (pooled from multiple cells) represented a spatial and temporal profile of the X-FP at SecGFP-GPI endocytic sites.

Using the pH pulsing assay, we found that the rise in pH 7 SecGFP-GPI intensity occurred only ∼3s prior to vesicle generation as opposed to nearly 40s for CME, followed by monitoring SecTfR internalisation (Ref. 5 & Figure S2c). Furthermore, we observed a poor correlation (r = 0.3) between pH 7 vs. 5 intensity per endosome, indicating that SecGFP-GPI is endocytosed without a major concentration in the endocytic vesicle (Figure 1c). In comparison to CME^5,27,28^, it should be noted that we faced two main challenges, i) lack of a cytoplasmic marker (for *e.g.* clathrin) for the endocytic site, ii) lack of strong concentration of the cargo prior to the endocytic vesicle pinching. Regardless, the protocol developed provided a reliable real-time assay for studying the spatio-temporal dynamics of the internalisation of GPI-anchored proteins (See, S.I.).

### Newly formed CG endocytic vesicles recruit GBF1, ARF1, and CDC42

We visualised the temporal dynamics of co-expressed CDC42 at the site of formation of nascent SecGFP-GPI endocytic vesicles. TagRFPt-CDC42 recruitment was quantified as average fold accumulation of CDC42 relative to the local background in all the endocytic events recorded over a 40s time window straddling the endocytic event. A significant change in the intensity of CDC42 began at around −9s and peaked around 0 to +3s (Figure S2a). Based on the presence or absence of a co-detected CDC42 spot (within −18 to +18s time window) at the endocytic site, we identified two populations via manual classification (See S.I. for a detailed description). We called them CDC42 Coloc (Co-detection of CDC42 and SecGFP-GPI) and CDC42 NoColc (the remainder). We compared the fold accumulation of all the CDC42 spots (CDC42 All), CDC42 Coloc and CDC42 NoColoc with Random. While the CDC42 Coloc profile was similar to that of CDC42 All, the CDC42 NoColoc profile was comparable to Random (Figure S2b). Thus, the endocytic sites detected by our assay consisted of two populations wherein one fraction exhibited an accumulation of CDC42 while the second fraction failed to show a discernable accumulation. CDC42 Coloc corresponded to the 56% (of the CDC42 All) SecGFP-GPI endocytic sites. As the removal of the events which did not coincide with the presence of CDC42 did not alter CDC42 recruitment profile (compare, Figure 1d & S1a), they were discarded from further analysis. While the reasons for not detecting CDC42 at all endocytic events is a function of both the signal and noise in the data, it may also reflect a genuine lack of recruitment at some endocytic events (See S.I. for a detailed explanation). Henceforth, for all X-FPs, we report pH pulsing traces that were classified as co-detected with the SecGFP-GPI (See S.I. and Table 2, for methods and statistical details and analysis).

**Table 1:** Correlation for indicated traces using Pearson’s correlation coefficient. n.s - not significant, p-value > 0.05 by Student’s t-test. ＠ - performed with one frame shift; # - performed for traces with spot radius = 2 pixels and background donut = 6-8 pixels. Rest of the calculations are done with traces with spot radius = 3 pixels and background donut = 6-8 pixels. 1 pixel = 84 nm.

| Molecule Pair | Time Interval (s) |  |
| --- | --- | --- |
|  | (-36s to -12s) | Specific interval |
| GBF1 vs CDC42 | 0.5767 |  |
| GBF1 vs LIFEACT | n.s. |  |
| GBF1 vs ARP3 | n.s. |  |
| GBF1 vs IRSp53 | n.s. |  |
| ARF1 vs GBF1 | 0.6323 |  |
| ARF1 vs CDC42 | 0.564 |  |
| ARF1 vs IRSp53 | n.s. |  |
| ARF1 vs ARP3 | 0.6347 |  |
| ARF1 vs LIFEACT | 0.7478 |  |
| CDC42 vs IRSp53 | 0.65<sup>#,@</sup> | 0.5698<sup>#</sup> (-33s to 0s) |
| CDC42 vs LIFEACT | 0.6087 | 0.54 (-36s to +6s) |
| CDC42 vs ARP3 | n.s. |  |
| ARP3 vs IRSp53 | n.s. | 0.7739<sup>#</sup> (-9s to +9s) |
| ARP3 vs LIFEACT | n.s. | 0.6292 (-33s to 0s) |
| ARF1 vs PICK1 | n.s. | 0.5698 (-36s to -12s) |
| PICK1 vs IRSp53 | -0.5353 |  |

n.s - not significant p-value considered here is 0.05 calculated via t-statistic.
@ - performed with one frame shift.

**Table 2:**
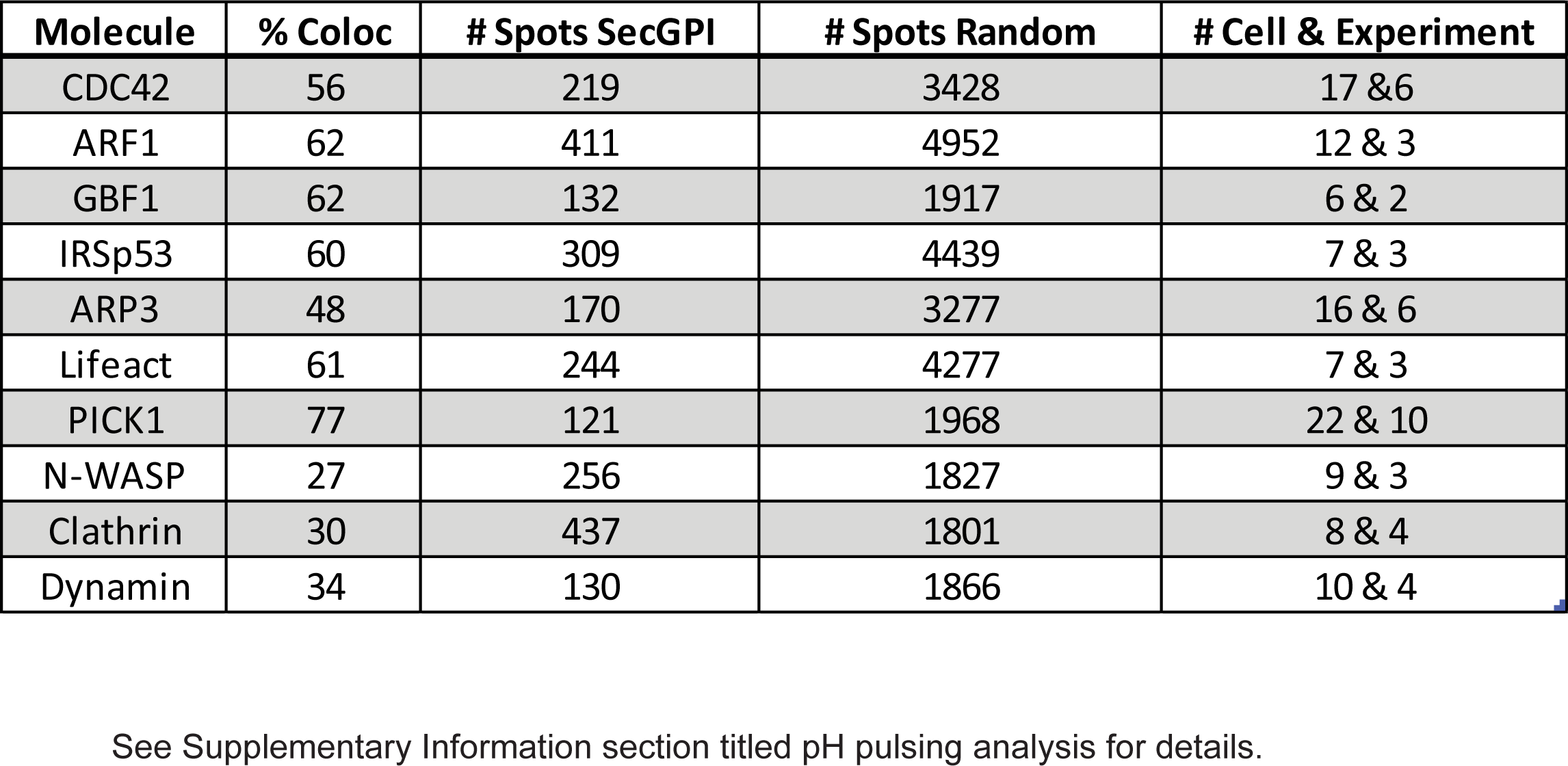
Details of pH pulsing assay data set. See S.I., section pH pulsing analysis for further details.

| **Molecule** | **% Coloc** | **# Spots SecGPI** | **# Spots Random** | **# Cell & Experiment** |
| --- | --- | --- | --- | --- |
| CDC42 | 56 | 219 | 3428 | 17 & 6 |
| ARF1 | 62 | 411 | 4952 | 12 & 3 |
| GBF1 | 62 | 132 | 1917 | 6 & 2 |
| IRSp53 | 60 | 309 | 4439 | 7 & 3 |
| ARP3 | 48 | 170 | 3277 | 16 & 6 |
| Lifeact | 61 | 244 | 4277 | 7 & 3 |
| PICK1 | 77 | 121 | 1968 | 22 & 10 |
| N-WASP | 27 | 256 | 1827 | 9 & 3 |
| Clathrin | 30 | 437 | 1801 | 8 & 4 |
| Dynamin | 34 | 130 | 1866 | 10 & 4 |

See Supplementary Information section titled pH pulsing analysis for details.

We next examined another CG pathway regulator, ARF1^15^ and found that mCherry-ARF1 was already accumulated at the forming CG endocytic sites at −36s and peaked at 0s (Figure 1e, S6a & Table 2). Predictably, mCherry-GBF1, an ARF1-GEF^17,33^ was also recruited to SecGFP-GPI spots (Figure 1f, S6b & Table 2). The temporal profile of GBF1 was correlated to ARF1 (r = 0.6, Table 1). When we extended the time window of observation for ARF1 and GBF1 further back in time (nearly 60s before scission), we concluded that the accumulation of ARF1 and GBF1 at CG endocytic sites inititates as early as −54s (Figure S2f-g). In absence of other molecules upstream of GBF1 and ARF1, this pair currently serves as the earliest initiators of the CG endocytic pathway. Furthermore, when we assessed the ultra-structure of newly formed fluid-filled endosomes by electron microscopy (EM)^18^ in cells treated with the small molecule inhibitor of GBF1, LG186^34^, the number of CLICs and fluid-uptake was drastically reduced, whereas, CME derived vesicles and uptake appear relatively unaffected (Figure S1e-f). In contrast to ARF1, GBF1 and CDC42, we did not observe a frequent recruitment of clathrin and dynamin to the CG endocytic sites. The recruitment profile was comparable to random for >65% endocytic events for both mCherry-Clathrin and Dynamin (Figure S2d-e & Table 2). The remainder of the profiles exhibited high levels of clathrin and dynamin at SecGFP-GPI sites in a fashion unlike that observed for CME (Ref. 5 & Figure S2c). This may account for the endocytic uptake of a fraction of SecGFP-GPI via a CME like process as shown previously (Ref. 12, Figure S1a-c) and might explain why we failed to quantitatively localise CDC42, ARF1 and GBF1 to all the SecGFP-GPI-endocytic events as noted above.

The pH pulsing analysis revealed a surprising facet of the recruitment of ARF1, which suggested a CDC42-independent function(s) for ARF1 in CG endocytosis. Despite being responsible for the timed removal of CDC42 from the plasma membrane^15^, ARF1 was recruited long before the recruitment of CDC42. Taken together these results establish a reliable real-time imaging assay to follow newly formed CG endocytic vesicles containing SecGFP-GPI, correlated with recruitment of its known regulators, ARF1, GBF1 and CDC42.

### Identification of BAR domain proteins involved in CG endocytosis

CLICs, visualised within 15s of their formation have a pleomorphic tubular appearance and lack a discernable protein coat when visualised by EM^18,22^. This prompted us to investigate the role of BAR domain proteins (BDPs) for their role in CG endocytosis due to their capability to sense or stabilise membrane curvature. Additionally, BDPs contain domains that can bind lipids and/or regulate actin machinery including RhoGTPases^29,30^. Thus, we performed a dsRNA screen in S2R+ cells^16,17^ to identify the BDPs involved in CG endocytosis using GBF1 (*garz*) and GFP dsRNA as positive and negative controls, respectively. The screen yielded 10 ‘hits’ which affected fluid-phase uptake (Figure 2a-b, S3). Predictably, Endophilin A required for dynamin-dependent CIE endocytosis of GPCRs and Shiga Toxin^10,11^ was not selected, whereas, GRAF1^31^ previously shown to be involved in CG endocytosis was identified as a ‘hit’. The other hits, Sorting nexin 6^35^ and Centaurinβ 1A^36^ have been implicated in late stages of endosomal trafficking; CIP4^37^ and NWK^38^ being dynamin interactors, were not pursued in this study. We focused, instead, on two classes of BDPs, MIM/CG32082 (I-BAR domain) and PICK1 (BAR domain), primarily due to their interactions with CDC42 and ARF1, respectively.

**Figure 2:**
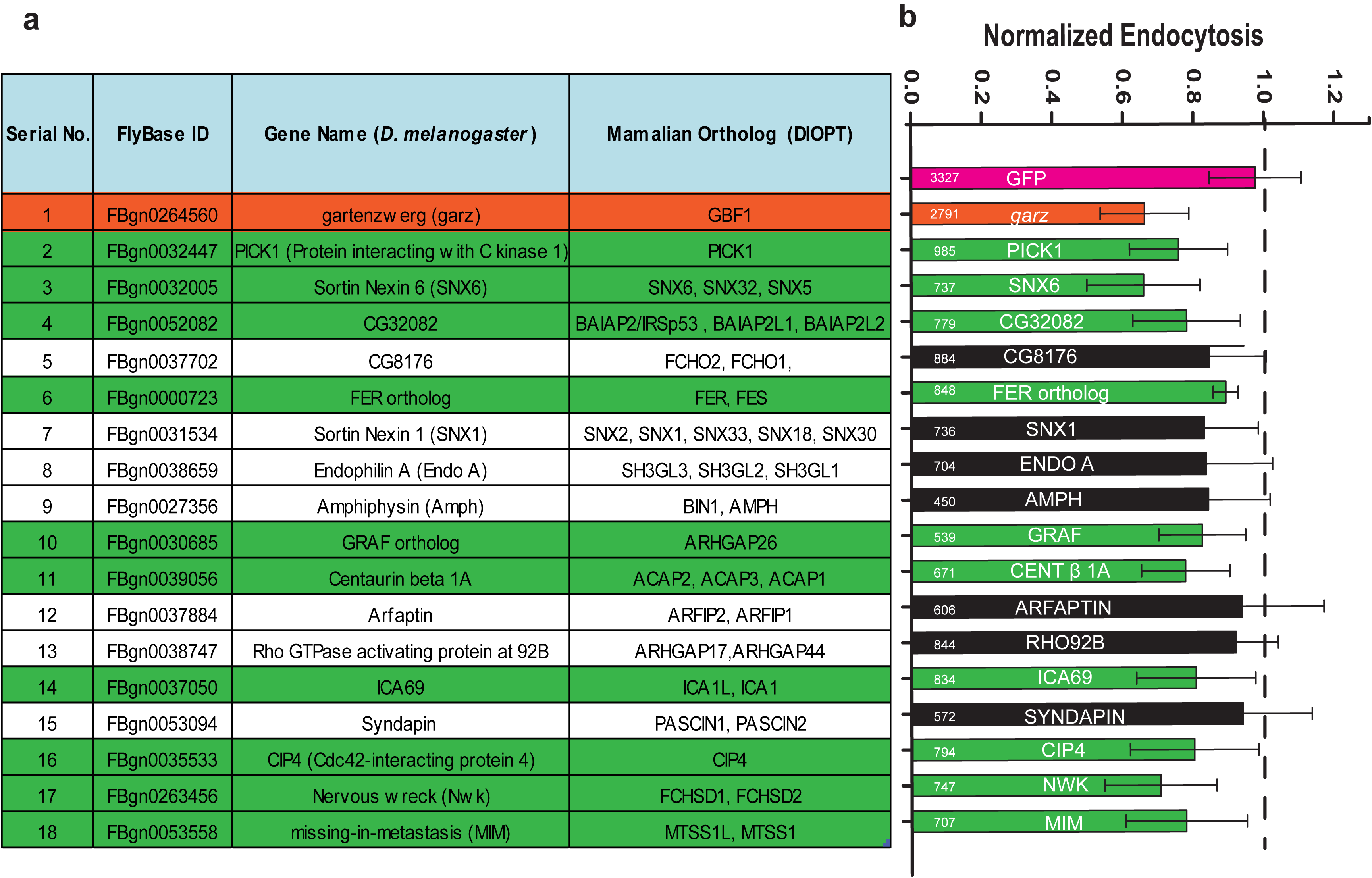
RNAi screen reveals BAR domain proteins involved in CG endocytosis. (**a**) List of *Drosophila* proteins in the PFAM database which contained one of the following BAR domains, PFAM IDs -PF06456, PF09325, PF06456, PF00611, and PF08397. The list was filtered to remove duplicates to give 21 genes. (**b**) Histogram shows normalized 5 minute fluid-phase uptake in S2R^+^ cells treated with 10ug of dsRNA for 4 days as indicated with dsRNA against GBF1 (*garz*) as positive, and GFP as negative controls. In a single experiment, mean uptake of one of GFP dsRNA coverslip was used to normalize the mean for rest of the coverslips. Data was pooled from 3 independent experiments and the cell numbers are indicated in the graph. The bars in green are significantly different from GFP dsRNA using 2-sample t-test (p < 0.05). PFAM database used: <u>ftp://ftp.ebi.ac.uk/pub/databases/Pfam/releases/Pfam27.0/</u>

### Role of the I-BAR domain containing IRSp53 in CG endocytosis

IRSp53, the mammalian orthologue of CG32082^39^, has been implicated in filopodia formation. IRSp53 has a multi-domain architecture including a CRIB, SH3 and PDZB domain and is known to interact with many actin regulatory proteins such as WASP-family verprolin-homologous protein 2 (WAVE2)^40–42^, Mena/VASP (vasodilator-stimulated phosphoprotein)^43,44^, Eps8^43,45–47^, mDia^41^ (Figure 3a). Furthermore, the recruitment of IRSp53 to the plasma membrane was compromised following ARF1 depletion^48^. Hence, IRSp53 was a good candidate to act as a signalling platform, linking CDC42 activation, membrane curvature and actin regulation for CG endocytosis.

**Figure 3:**
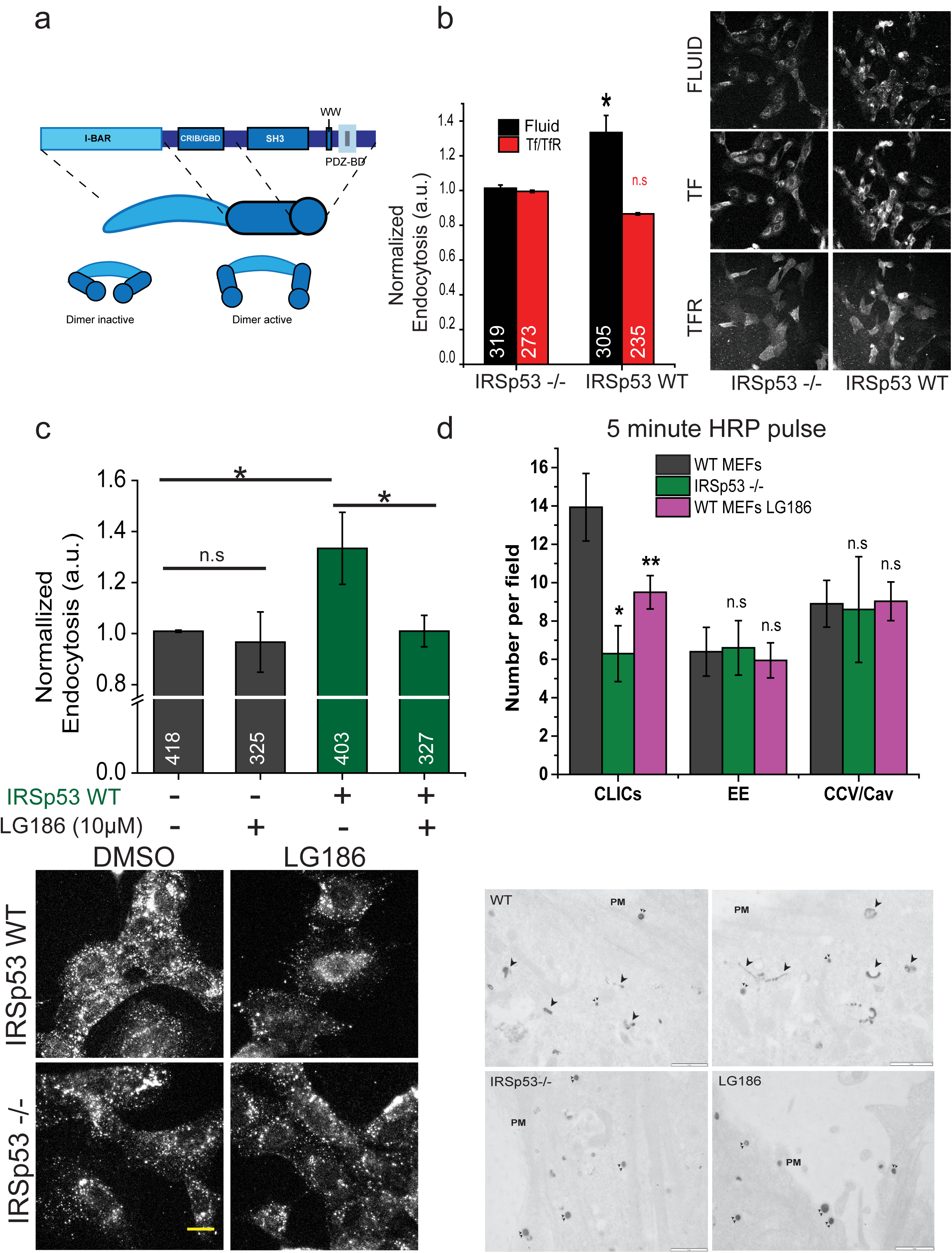
IRSp53 is involved in CG endocytosis. (**a**) Schematic depicting the domain organization of IRSp53. IRSp53 exists in an inactive dimer state, which upon binding to GTP-CDC42 gets activated, allowing SH3 domain to bind to its effectors. Histogram (left) shows 5 minute uptake of TfR and fluid-phase in IRSp53 WT cells normalized to IRSp53 -/- cells, along with representative images (right). Data is pooled from 2 independent experiments and the number of cells indicated in the figure. (**c**) Histogram (top) shows normalized 5 minute fluid-uptake in IRSp53-/- and IRSp53WT cells when treated with LG186 or vehicle (DMSO) along with representative images (bottom). Data is pooled from 2 independent experiments and the number of cells indicated in the figure. (**d**) Histogram (top) shows average number of endocytic structures quantified per field from the electron microscope images (bottom). Data pooled from 3 independent blocks. Untreated WT MEFs (WT, top row), IRSp53 null MEFs (IRSp53-/-, bottom left) or LG186-treated WT MEFs (LG186, bottom right) were incubated for 5 minute at 37°C with 10mg/ml HRP as a fluid phase marker before processing for electron microscopy. Endocytic structures close to the plasma membrane (PM) are filled with the electron dense peroxidase precipitate. WT cells show a range of endocytic structures including vesicular structures (double arrowheads) and tubular/ring-shaped putative CLIC/GEECs (large arrowheads) but the IRSp53-/- cells and LG186-treated cells only show predominant labeling of vesicular profiles. *p-value* < 0.05 (∗), 0.001 (∗∗) Mann Whitney U test (**b-c**) and 2-sample student’s t-test (**d**). Error bars, s.d (**b-d**) Scale bar, 20µm (**b-c**), 1µm (**d**) respectively. Schematic (**a**) was adapted with permission from MBInfo (<u>www.mechanobio.info</u>) Mechanobiology Institute, National University of Singapore.

To address the function of IRSp53 we compared endocytosis between mouse embryonic fibroblasts (MEFs) generated from IRSp53-/- mice (IRSp53-/- MEFs) and IRSp53-/- IRSp53WT add back MEFs (IRSp53WT MEFs)^43^. Loss of IRSp53 caused a significant reduction in fluid-phase uptake, without affected TfR internalisation (Figure 3b). We next addressed the nature of endocytosis in IRSp53 -/- MEFs. Fluid-phase uptake in IRSp53-/- MEFs remained refractory to LG186 mediated GBF1 inhibition (Figure 3c). By contrast, GBF1 inhibition in IRSp53WT MEFs decreased fluid-phase uptake to the levels observed in IRSp53-/- MEFs (Figure 3c) while endocytosed TfR remained unaffected (Figure S4a). We confirmed the lack of the CG endocytic pathway in IRSp53-/- MEFs by ultrastructural analysis of endocytic structures marked by the fluid-phase marker, HRP using EM^15,18^. We observed a significant reduction of CLICs in IRSp53-/- MEFs when compared with WT MEFs, while the number of clathrin and caveolae derived structures was relatively unaffected. Similar to Figure S1g, the CLICs are reduced significantly upon LG186 treatment in WT MEFs as well (Figure 3d, S4b).

This led us to hypothesise that CG cargo would traffic via CME in the absence of IRSp53. Therefore, we looked at the trafficking of GPI-AP (GFP-GPI), fluid-phase and TfR (CME) in the absence of IRSp53 at a higher resolution. We first, we counted the number of GFP-GPI and fluid endosomes and found them to be significantly lower in IRSp53-/- MEFs relative to IRSp53WT MEFs (Figure 4a). Conversely, TfR endosomal number was unaffected (Figure 4a). We next, looked at co-localisation of GPI-AP/fluid-phase with TfR. A relatively higher fraction of GFP-GPI (Figure 4b) and fluid-phase co-localized with co-pulsed Tf (Figure 4c) in IRSp53-/- MEFs than IRSp53WT MEFs. Thus, removal of IRSp53 specifically affected fluid-phase and GPI-AP endocytosis, while CME remained unaffected. Moreover, in cells lacking IRSp53, GPI-AP and the residual fluid-phase is endocytosed via CME.

**Figure 4:**
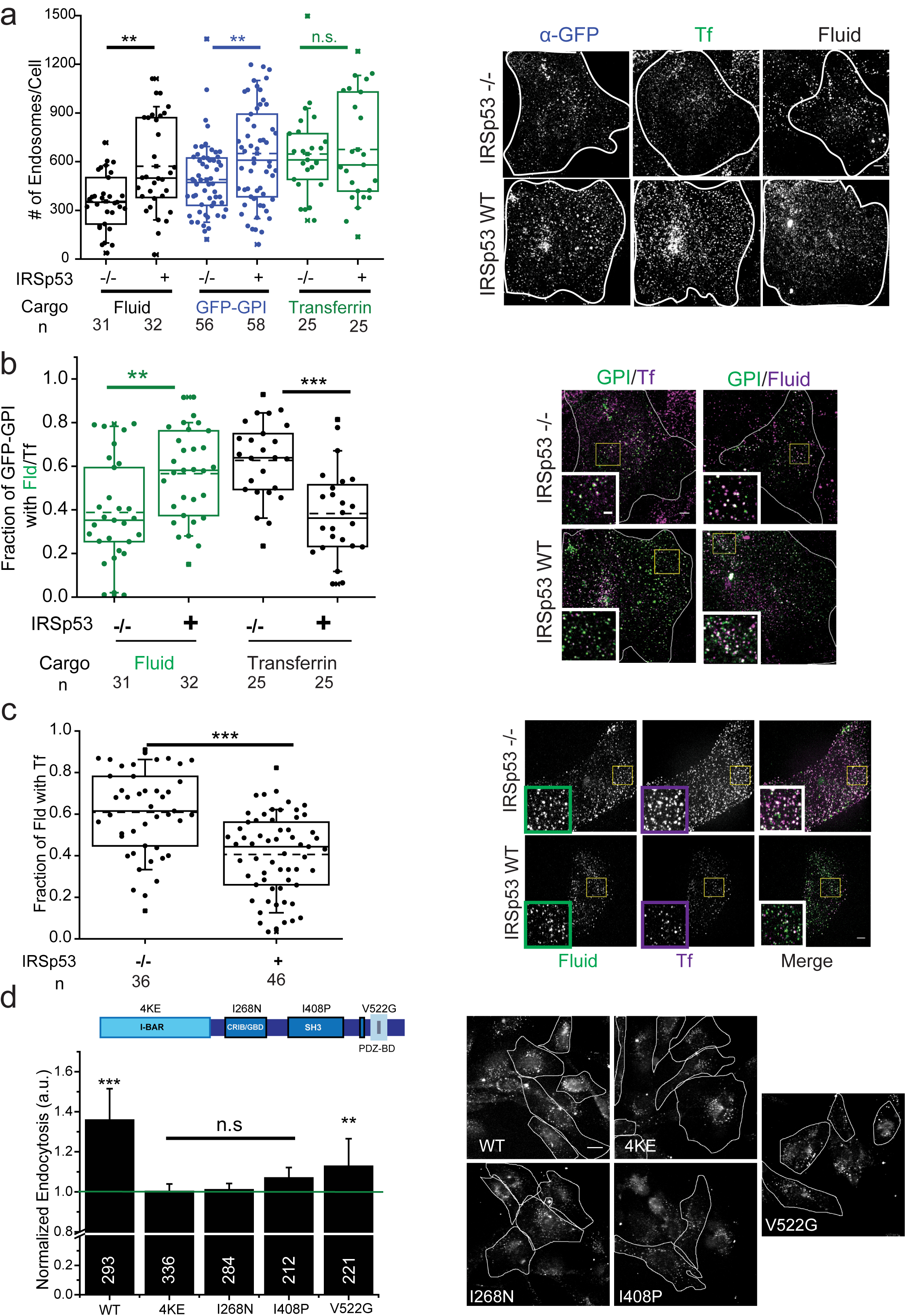
CG pathway is abolished in the absence of IRSp53. (**a**) Box plot shows the number of endosomes per cells (left) for endocytosed GFP-GPI (anti-GFP Fab), fluid-phase and TfR in IRSp53 -/- and IRSp53 WT cells when pulsed for 2 minutes along with representative images (right). Data is pooled from 2 independent experiments and the number of cells indicated below the graph. (**b**) Plot (left) shows quantification of the fraction of GFP-GPI endocytic vesicles containing fluid-phase or Tf. Images (right) show representative single confocal slices of a 2 minute pulse of anti-GFP Fab (green)/TMR-Dextran (magenta) and anti-GFP Fab (green)/A568-Tf (magenta) in IRSp53-/- (top row) and IRSp53WT (bottom row) cells. Inset depicts magnified views of the indicated region; single channel images are in panel 4a. Data is pooled from 2 independent experiments and the number of cells is indicated below the graph. (**c**) Plot (left) showing quantification of the fraction of 1-minute fluid-phase endocytic vesicles containing Tf. Images show representative single confocal slices of a 1 minute pulse of TMR-Dextran (green) and A647-Tf (magenta) in IRSp53-/- (top row) and IRSp53WT (bottom row) cells. Inset depict magnified views of the indicated region. Data is pooled from 2 independent experiments and the number of cells indicated below the graph. (**d**) Histogram (left) shows 5-minute uptake of fluid-phase in IRSp53 -/- MEFs transduced with virus expressing GFP-IRSp53 WT, GFP-IRSp53 4KE, GFP-IRSp53 I268N, GFP-IRSp53 I408P and GFP-IRSp53 V522G, normalized to that in IRSp53 -/- MEFs, along with representative images (right). Data is pooled from 2 independent experiments and the number of cells indicated in figure except for IRSp53-/- (381). *p-value* < 0.01 (∗), and 0.001(∗∗) by Mann-Whitney U Test (**a-d**). Scale bar, 20µm (**d**), 5µm (**a-c**) respectively. Error bars, (**d**) represent s.d.

We next analysed the contribution of different domains of IRSp53 on CG endocytosis by re-introducing into IRSp53-/- MEFs, GFP-tagged IRSp53WT and a number of mutants of IRSp53 specifically defective in various domains (Figure 4d, schematic). We found that GFP-IRSp53WT & GFP-IRSp53V522G rescued endocytosis while the rest of the mutants failed to do so (Figure 4d). In conclusion, these results indicated that IRSp53 is an essential and specific regulator of CG endocytosis that requires functional I-BAR, CRIB and SH3 domains.

### IRSp53 is recruited to forming CG endocytic vesicles

The complete absence of CG endocytosis in IRSp53-/- led us to hypothesise that IRSp53 has a direct role to play in CG vesicle formation. Hence, we used pH pulsing assay and examined the recruitment of mCherry-IRSp53 to the forming SecGFP-GPI endocytic sites. A majority of (>60%) endocytic events exhibited prominent recruitment of IRSp53 (Figure 5a, S6c & Table 2). Since the I-BAR domain of IRSp53 has been shown to sense/induce negative curvature in a membrane tension and protein concentration dependent manner^49^, we looked at changes in its spatial distribution during the formation of endocytic vesicle using two types of masks – a spot and a ring mask (Figure 5a, schematic). Unlike the intensity profiles of CDC42 that did not exhibit any differential temporal patterns of recruitment between the two types of masks (Figure 5b, black vs. red trace), IRSp53 displayed a biphasic recruitment pattern (Figure 5a). In phase I (−36 to −15s), IRSp53 was first recruited over a large area indicated by an increase in intensity in both spot and ring traces. Subsequently, in phase II (−15 to 0s), IRSp53 was confined at the centre indicated by a concerted decrease in the ring mask intensity and increase in the spot mask intensity trace (Figure 5a, black vs. red trace). The increase of IRSp53 was more prominent in phase II toward the centre (Figure 5a, black vs. purple trace) and correlated with the recruitment of CDC42 (r = 0.6; Table 1).

**Figure 5:**
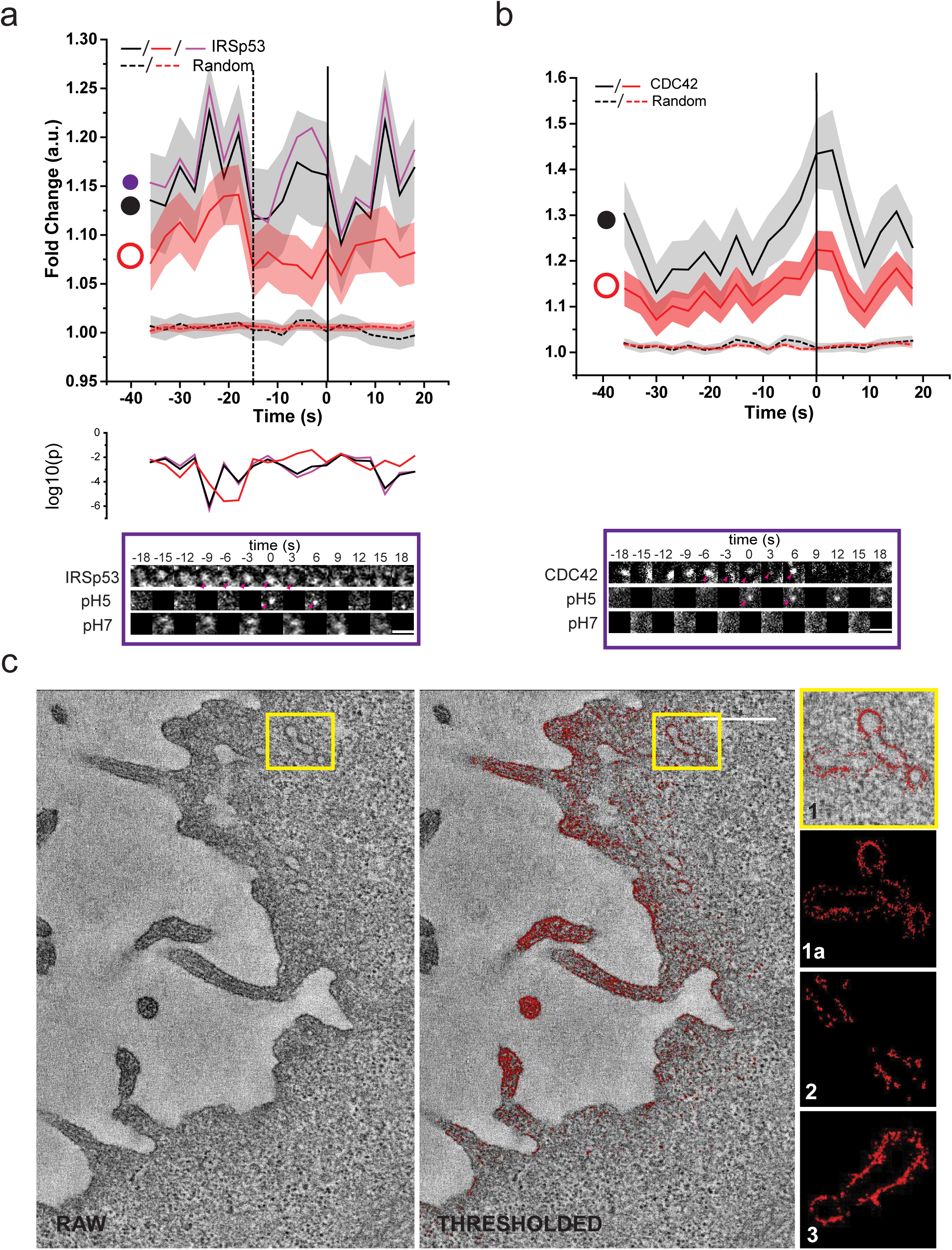
IRSp53 is recruited to forming CG endosomes. (**a**) Graphs show the average normalized fluorescence intensity versus time traces for the recruitment of three different regions [circles, violet, r = 170 nm and black (r = 250 nm) and annulus, orange (r = 250-420 nm)] for the recruitment of IRSp53-mCherry to the forming SecGFP-GPI endocytic sites and its corresponding random intensity trace (n, Table2). (**b**) Graphs show the average normalized fluorescence intensity versus time traces for the recruitment of TagRFPt-CDC42 to the forming SecGFP-GPI endocytic sites and its corresponding random intensity trace to two different regions [circle, black, r = 250 nm; and annulus, orange (r = 250-420 nm)]. Error bars, (**a-b**) represent s.e.m (n, Table 2). The random traces were derived from randomly assigned spots of the same radius as the endocytic regions, as detailed in S.I. Endocytic distribution at each time point was compared to the random distribution (**a**) by Mann-Whitney U test and the log_10_ (p) is plotted below each trace [log_10_ (0.05) is −1.3 and log_10_ (0.001) is −2.5]. Representative montages is depicted below the graphs (**a-b**). Arrowheads indicate the newly formed endocytic vesicle. (**c**) Electron micrographs of AGS cells cotransfected-GFP-IRSp53 and GFP binding protein coupled to Apex (GBP-Apex). The DAB reaction was performed and the cells were processed for electron tomography (See S.I.). A single section of the original tomogram (left) and density-based thresholded of the same plane (middle) reveal electron dense structures containing IRSp53 at membrane surfaces. The whole of PM of the tomographic volume was rendered and different examples of enlarged tubular regions of interest show GFP-IRSp53 recruitment patterns (right). Scale bar, 1.5µm (**a-b**) and 0.5µm (**c**) respectively.

To visualise the recruitment of GFP-IRSp53 at high spatial resolution, we coexpressed a GBP-APEX reagent (GFP binding protein-soybean ascorbate peroxidase) and processed for EM as described previously^50^ (Figure 5c, S4c, Supplementary Movies 3-4). GBP-APEX binds to GFP and converts 3,3’-diamino-benzamidine into an osmiophilic polymer in presence of H_2_O_2_^50^. Images of 3D rendering from the electron densities revealed that IRSp53 associated with tubular structures characteristic of CLICs as described previously^18^ and as expected, with filopodial structures^39^ (Figure 5c, See SI). In the 2D sections, IRSp53 was observed to accumulate as discrete patches at the plasma membrane (PM) (Figure S4c i-ii, arrowheads) and was frequently associated with tubulovesicular invaginations or tubular structures close to PM and filopodial tips (Figure S4c iii, double arrowheads).

### ARP2/3 based F-actin polymerization is necessary for CG endocytosis

A functional CG endocytic pathway requires dynamic actin since inhibition of actin polymerization (latrunculin A), or filament stabilisation (jasplakinolide), inhibited CG endocytosis^14^. CDC42 and IRSp53 are a core component of a signalling axis that indirectly controls the location and activity of the ARP2/3 actin nucleation complex^40,51,52^ (Figure S4d). More relevantly, CK666^53^ mediated inhibition of ARP2/3 complex, impaired both fluid-phase and TfR uptake in a dose dependent manner. However, the extent of inhibition of CG internalisation was markedly more prominent (Figure 6a). By contrast inhibition of formins via SMIFH2^54^, failed to inhibit CG endocytosis (Figure S5a).

**Figure 6:**
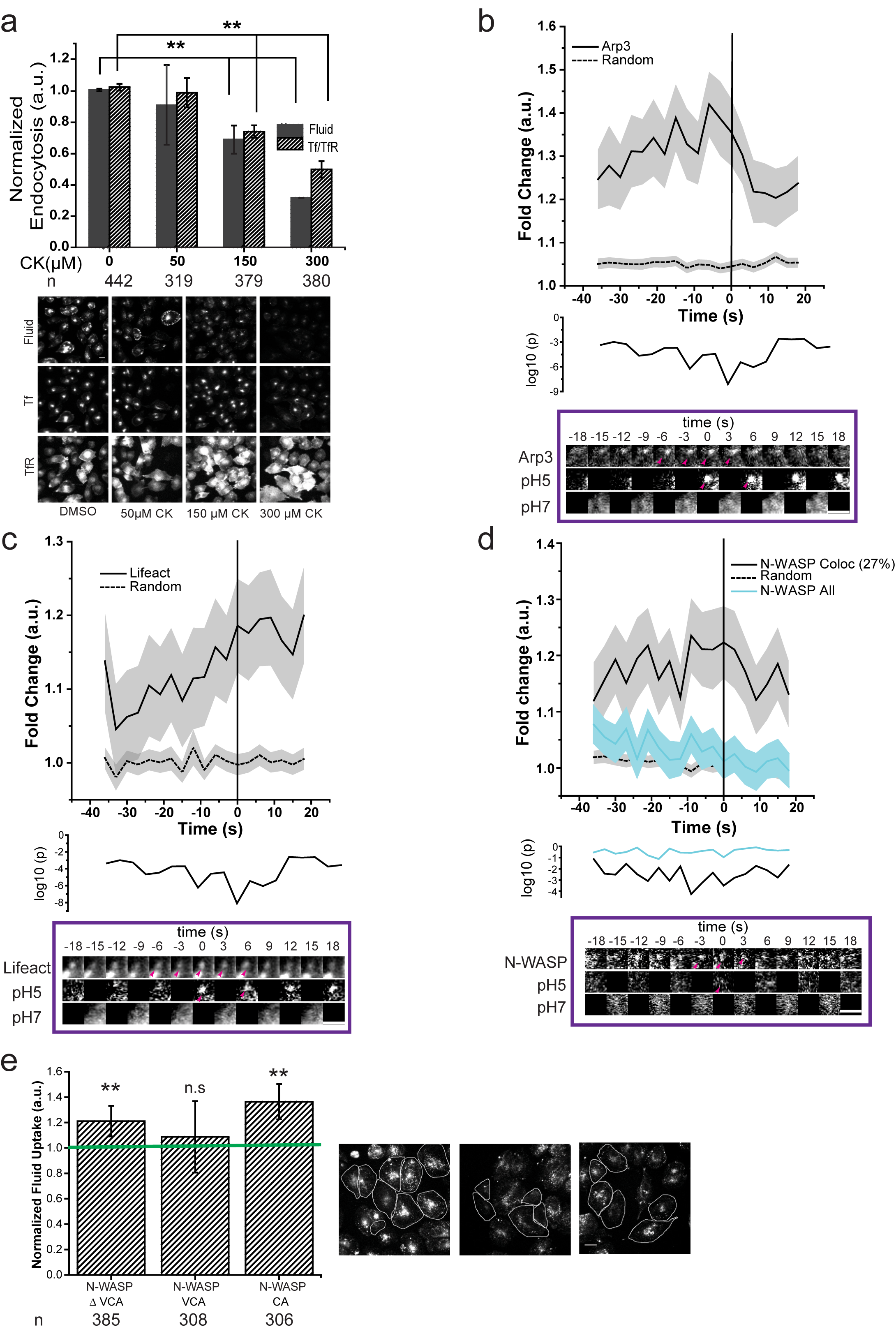
Arp2/3 based actin machinery is required for CG endocytosis. (**a**) Histograms (top) shows quantification of fluid-phase and TfR uptake in AGS cells treated with DMSO alone (0µM) or the indicated concentrations of ARP2/3 inhibitor, CK666, normalized to DMSO treated controls, along with its representative images (below). Data is pooled from 2 independent experiments and the number of cells shown indicated the graph. (**b-d**) Graphs show the average normalized fluorescence intensity versus time traces for the recruitment of mCherry-ARP3 (**b**), pRuby-Lifeact (**c**) and mCherry-NWASP (**d**) to the forming SecGFP-GPI endocytic sites, and its corresponding random intensity trace (n, Table 2). The random traces were derived from randomly assigned spots of the same radius as the endocytic regions, as detailed in S.I. Endocytic distribution at each time point was compared to the random distribution by Mann-Whitney U test and the log_10_ (p) [log_10_ (0.05) is −1.3 and log_10_ (0.001) is −2.5] is plotted below each trace (**b-d)**. Representative montages are depicted below the graphs. Arrowheads indicate the newly formed endocytic vesicle. (**e**) Histogram (left) shows normalized 5 minute mean fluid-phase uptake in AGS cells over-expressing pIRESCA domain, GFP-VCA domain and GFP-N-WASP∆VCA from N-WASP compared to untransfected cells and representative images (right). The transfected cells are outlined. Data is pooled from 2 independent experiments and the number of cells shown below the graph. Error bars represent s.e.m. (**b-d**) and s.d. (**a** & **e**). *p-value* < 0.01 (∗), and 0.001(∗∗) by Mann-Whitney U Test (**a** & **e**). Scale bar 1.5µm (**b-d**), 20µm (**a** & **e**).

We next explored the spatio-temporal dynamics of F-actin and ARP2/3 complex using pRuby-lifeact^55^ and mCherry-ARP3 respectively during the formation of the endocytic vesicle. ARP3 recruitment (Figure 6b, S6d & Table 2) began earlier than −35s and peaked at −6s. This was unexpected since CDC42, a key regulator of ARP2/3^56,57^ was not recruited until −9s. Instead, the ARP3 profile was correlated with IRSp53 between −9s and +9s (r = 0.8; Table 1). This indicated that, at least initially, the ARP2/3 complex may be recruited in a CDC42-independent manner. On the other hand, F-actin recruitment began around −9s and continued even after the scission event (Figure 6c, S6e & Table 2), highly correlated to the CDC42 profile (r = 0.6; Table 1). Thus, F-actin is generated coincident with the recruitment of CDC42, a known regulator of actin polymerization^56,57^. These observations suggest that ARP2/3 complex might be first recruited as an inactive ARP2/3 complex, and then activated following the arrival of CDC42.

### PICK1, a negative regulator of the ARP2/3 complex, is recruited to forming CG endosomes

To address how ARP2/3 may be maintained at the forming endocytic pit in an inactive state we analysed the role of PICK1, another ‘hit’ in the screen (Figure 2). PICK1, a highly conserved protein, possesses PDZ and BAR domain (Figure 7a), that by intramolecular interaction, maintain PICK1 in an auto-inhibited state. This auto-inhibited state is further stabilised upon GTP-ARF1 binding to the PDZ domain^58^. Additionally, activated PICK1 negatively regulates ARP2/3 mediated actin polymerization^58–60^. The ability of PICK1 to inhibit ARP2/3 complex is suppressed by GTP-ARF1^58^. To confirm a role of PICK1 in CG endocytosis in mammalian cells, we utilised a specific small molecule inhibitor of PICK1, FSC231^61^. CG endocytosis (fluid-phase) was inhibited in a dose-dependent fashion (Figure 7b). Additionally, in stable PICK1 knockdown lines, fluid-phase and folate receptor (FR-GPI, another GPI-AP) endosomal number were lower than that measured in scrambled shRNA stable lines, while TfR endocytosis remained unaffected (Figure 7c-d).

**Figure 7:**
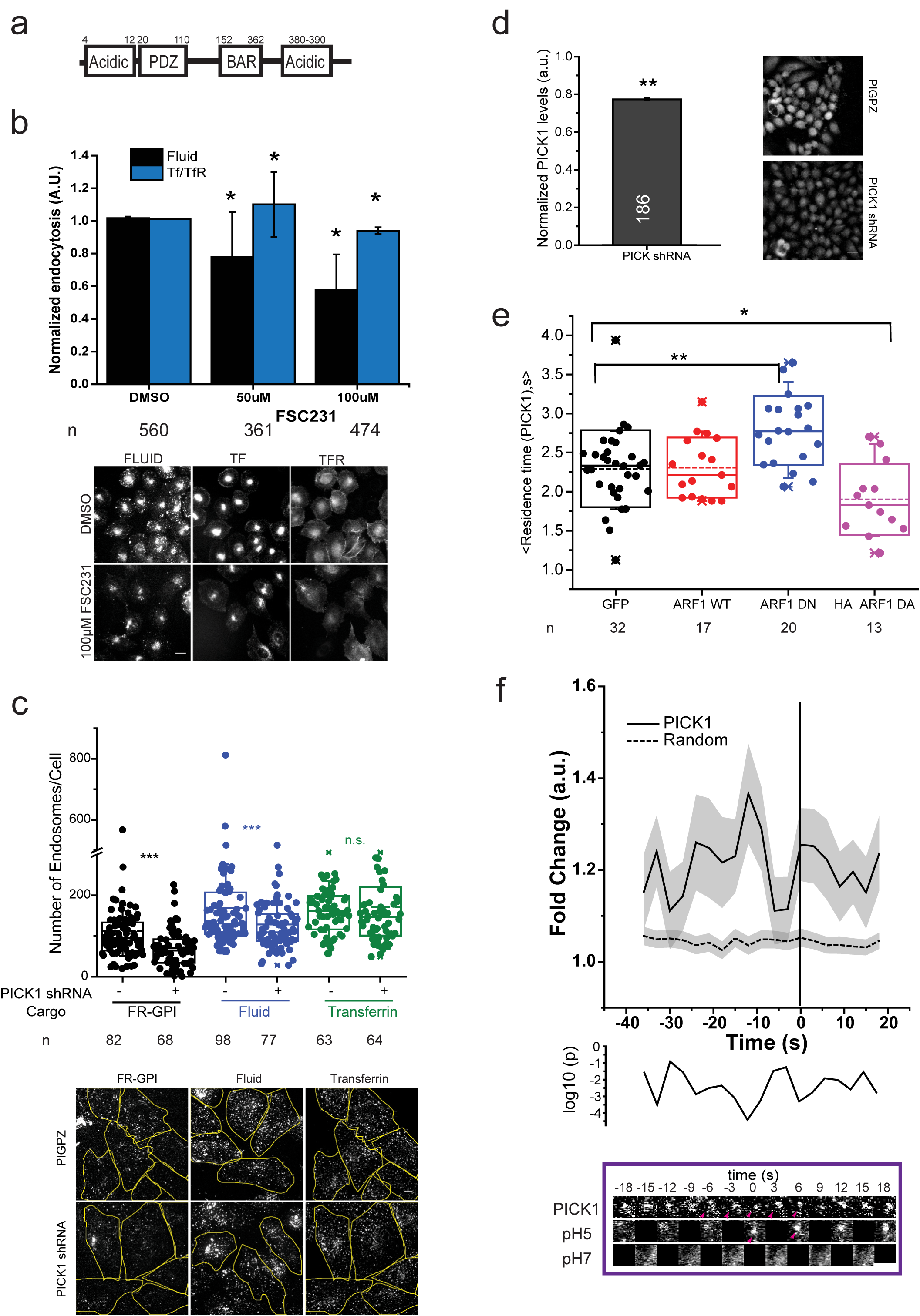
PICK1 is involved in CG endocytosis and is negatively regulated by ARF1. (**a**) Schematic depicts domain organization of PICK1. (**b**) Histograms (top) shows quantification of fluid-phase and TfR uptake in AGS cells treated with DMSO alone (0µM) or the indicated concentrations of PICK1 inhibitor, FSC231, normalized to DMSO treated controls, along with its representative images (below). Data is pooled from 2 independent experiments with number of cells shown below the graph. (**c**) Box plot (top) shows the number of endosomes per cells for FR-GPI (Cy3-Mov18), fluid-phase and TfR in scrambled (PIGPZ) and PICK1 shRNA infected AGS cells when pulsed for 2 minutes along with representative images (bottom). Data is pooled from 2 independent experiments and the number of cells indicated below the graph. (**d**) Histogram (left) shows normalized PICK1 levels measured by immunostaining in PICK1 shRNA infected AGS cells along with representative images (right). Data is pooled from 2 independent experiments with number of cells indicated in the figure except PIGPZ (292). (**e**) Box plot (top) shows the residence time of TagRFPt-PICK1 spots at the TIRF plane (See S.I.), averaged in an individual cell expressing either GFP, GFP-ARF1 WT, GFP-ARF1 DN or HA-ARF1 DA. The data are pooled from 2 independent experiments with cell number indicated below the graph. (**f**) Graph shows the average normalized fluorescence intensity versus time trace for the recruitment of TagRFPt-PICK1 to the forming SecGFP-GPI endocytic sites and its corresponding random intensity trace (n, Table 2). The random traces were derived from randomly assigned spots of the same radius as the endocytic regions, as detailed in S.I. Endocytic distribution at each time point was compared to the random distribution by Mann-Whitney U test and the log_10_ (p) [log_10_ (0.05) is −1.3 and log_10_ (0.001) is −2.5] is plotted below. Representative montage is depicted below. Arrowheads indicate the newly formed endocytic vesicle. Error bars represent s.e.m. (**f**) and s.d. (**b** & **d**). *p-value* < 0.01 (∗), 0.001(∗∗) and 0.0001 (∗∗∗) by Mann-Whitney U Test (**b-e**). Scale bar 1.5µm (**f**), 20µm (**b** & **d**) and 5 µm (**c**).

Predictably, GFP-ARF1 and TagRFP-PICK1 co-localized in punctate spots at the TIRF plane in accordance with previous reports^58^ (Figure S5b). To test the effect of ARF1 activity on PICK1 recruitment we co-expressed TagRFP-PICK1 with either ARF1-WT or dominant negative (ARF1-DN; T31N) and active mutant (ARF1-DA; Q71L) and tracked the number and dynamics of PICK1 spots using TIRF microscopy (See, S.I.). The residence time of PICK1 increased significantly in the presence of ARF1-DN while it was reduced in presence of ARF1-DA (Figure 7e). ARF1-DN and DA have been shown to decrease and increase endocytosis respectively^15^. Thus, the local presence of ARF1-GTP resulted in the removal of PICK1 from the membrane. Thus, we hypothesised that PICK1 was recruited to forming CG endosome at the early stage, rendering ARP2/3 inactive.

To explore this possibility, we utilised the pH pulsing assay to directly visualise PICK1 at the CG endocytic site. We found that TagRFP-PICK1 was recruited to forming CG endocytic sites (Figure 7f, S6f & Table 2) in a pulsatile fashion. Maximum enrichment occurred at −12s, with an eventual loss corresponding to the time of the rapid rise in ARF1 recruitment around −9s (Figure 1e, r = 0.6, Table 1). Thus, pH pulsing assay led to the realisation that in CG endocytosis, the interplay of two BDPs, PICK1 and IRSp53 regulate ARP2/3 complex. These BDPs interact with the ARP2/3 complex (Figure S5b, upper panel and Figure S4d), regulate its activity at the forming CG endocytic sites, in opposing fashion under the influence of ARF1 and CDC42.

## DISCUSSION

CG endocytosis was initially discovered as a route for the entry of toxins^62^, and fluid-phase and GPI-anchored proteins when CME^12,13,63^ was perturbed, raising some concerns regarding its physiological role in unperturbed cells^19^. Using pH pulsing assay we show, here, that a majority of SecGFP-GPI-containing endocytic vesicles form due to a stereotypical and temporally orchestrated recruitment of the key molecular machinery namely, CDC42, ARF1 and GBF1, which, in turn, mediate the coordinated assembly of specific BAR-containing, membrane deforming and actin regulatory proteins, IRSp53 and PICK1. Thus, a dedicated complex protein machinery drives CG internalisation, similarly to what is observed in CME. Notably, however, the vast majority of the endocytic vesicles are devoid of clathrin and dynamin. The ability of the pH pulsing assay to provide a temporal profile for the recruitment dynamics of the molecular players has considerably extended our understanding of CG endocytic vesicle formation.

The GBF1/ARF1 pair is the earliest module that is assembled and judging by their recruitment profiles, it takes around 1 minute and more, to assemble the molecular machinery for CG endocytosis. How this pair is concentrated at a forming CG endocytic site is an open question. The CG machinery includes CDC42, ARP2/3 and F-actin along with BDPs, PICK1 and IRSp53, and in the model (Figure 8) we propose a biphasic mechanism. ARF1 recruitment occurs roughly in two phases. In the first phase the accumulation is slow, accompanied by the presence of PICK1 and absence of CDC42. In the second phase, beginning around — 9s (before scission), the accumulation of ARF1 speeds up concomitant with the arrival of CDC42 and loss of PICK1. How the kinetics of ARF1 recruitment is regulated is unclear since its GAP (GTPase activating protein) is presently unknown, as is the GEF for CDC42. The presence of PICK1 provides an explanation of how ARP2/3 is recruited to the forming CG endocytic vesicles long before CDC42, and is maintained in an inactive state. ARP2/3 is then induced to promote actin branching only upon the arrival of IRSp53 and CDC42.

**Figure 8:**
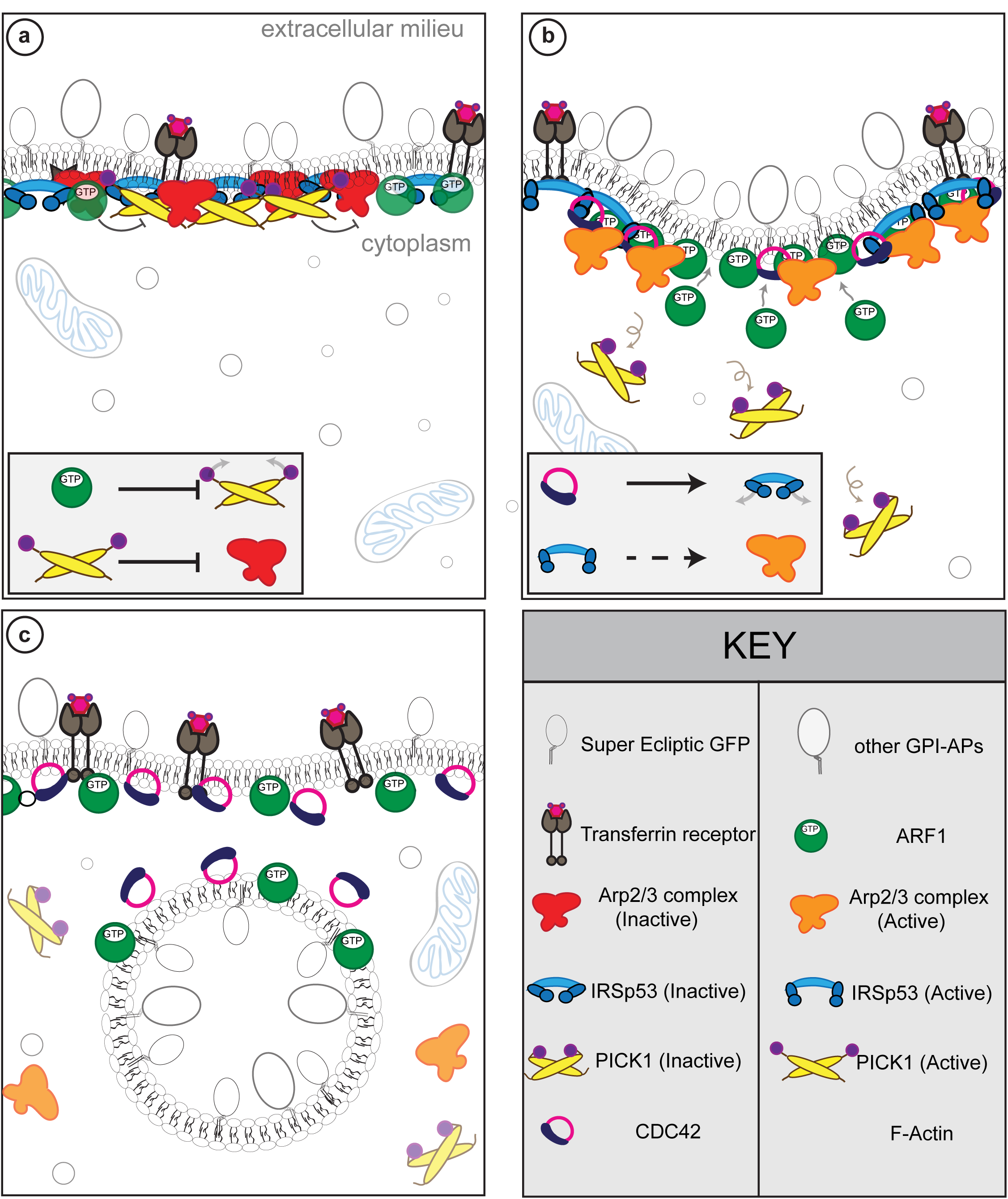
Schematic depicting the proposed biphasic mechanism for CG endocytic vesicle formation. (**a**) Phase Ι: Characterized by recruitment of ARF1/GBF1, PICK1, ARP2/3 and IRSp53 but not the buildup of F-actin and CDC42. Here IRSp53 may be recruited by its I-BAR domain in the absence of GTP-CDC42, keeping its SH3 domain in an intra-molecular inhibited state. PICK1 keeps ARP2/3 in an inactive state. (**b**) Phase ΙΙ: Characterized by recruitment of CDC42, sharp increase in ARF1 leading to removal of PICK1. This allows the activation of ARP2/3 and buildup of F-actin. CDC42 binds to the CRIB domain of IRSp53 thereby activating it. The SH3 domain of IRSp53 can now bind to ARP2/3 activators and create F-actin. (**c**) Phase III: Characterized by endocytic vesicle formation, presence of CDC42, ARF1/GBF1, and F-actin.

The role of ARP2/3 in CG endocytosis is reminiscent of the endocytic process occurring in the budding yeast. In this system, the endocytic machinery strictly depends on Las17, the yeast homologue of N-WASP but not so much on clathrin and dynamin^64,65^. There are however important differences. In CG endocytosis, the ARP2/3 complex appears to be activated independent of its canonical NPF, N-WASP, a CDC42 effector^57^. N-WASP neither showed any recruitment to the majority of the forming SecGFP-GPI spots (Figure 6d and S6g) nor did the over-expression of its dominant negative mutant affect fluid-phase endocytosis (Figure 6e). By contrast, in CME, both ARP2/3 complex and N-WASP are recruited to budding CME vesicles, and influence endocytosis in some cell types^27^.

The unexpected recruitment profile of ARP2/3, and the identification of two BDPs, PICK1 and IRSp53 as upstream regulators of ARP2/3 activity also suggests a biphasic mechanism. PICK1 operates in the early phases, and may function as an inhibitor of ARP2/3, consistent with the modest recruitment of F-actin in the presence of PICK1 observed here and as reported previously^58,59^. PICK1 recruitment occurs via its BAR domain since when mutated and overexpressed, it acts as a dominant negative for CG endocytosis (Figure S5c). PICK1 recruitment is not only rapid but also transient. We find that the localisation of PICK1 to the membrane was negatively correlated to the activity of ARF1, similar to that observed in neuronal cells^58,66^ wherein GTP-ARF1 interaction with PICK1 rendered PICK1 incapable of inhibiting ARP2/3 complex. This is followed by the second phase characterised by the simultaneous recruitment of CDC42/IRSp53 effector complex enabling the activation of ARP2/3 and subsequent polymerization of actin at the site of endocytosis. However, the NPF linking IRSp53 and ARP2/3 activation is not yet characterised and is the subject of investigation.

There is no single unifying theme for vesicle scission in CIE and multiple modules may co-exist. Recently, Endophilin A has been shown to facilitate tubule scission by a combination of scaffolding, dynamin recruitment, and dynein-mediated elongation of membrane tubules leading to an increase in friction^67^. In the CG pathway, IRSp53 emerges as a major player. This protein may function by multiple mechanisms: it can couple negative curvature with membrane tension, it can scaffold membrane at moderate densities^49^, it can regulate actin polymerization as it does in filopodia formation^39^. A minimal model accounting for all these activities suggests that IRSp53 might be enriched at the vesicle neck, where it would regulate the actin machinery necessary to trigger CG vesicle scission. The spatio-temporal dynamics of IRSp53 at CG sites in consistent with this model. Furthermore, the complete and specific loss of CG endocytosis (but not CME) in the absence of IRSp53 makes IRSp53 essential for CG endocytic process.

In summary, we propose that CG endocytic vesicle formation begins with GBF1/ARF1 concentrating at sites of endocytic pits. Following this, initial event ARP2/3 is recruited but is held in an inactive state by PICK1 (Figure 8a). Meanwhile, IRSp53 is recruited (potentially via its I-BAR domain) and activated by CDC42 leading to ARP2/3 activation via unknown effector(s) (Figure 8b). The loss of IRSp53 and ARP2/3 from the membrane as the endocytic vesicle is pinched is consistent with their role in endosomal neck dynamics, providing a new candidate for molecular machinery of the pinching process in the absence of dynamin. It is conceivable that the IRSp53 provides a scaffold for a friction-based scission mechanism as recently suggested^67^, with actin polymerization providing driving force for tubule elongation. Alternatively, this force could arise from the involvement of a microtubule-based machinery as recently advocated in the internalization of cholera toxin ^68^. The assays developed here and the identification of a number of molecular players and their temporal recruitment profile provides a path towards understanding a molecular mechanism for the formation of a CG endocytic vesicle.

## METHODS

### Cell culture, reagents, and plasmids

See Supplementary Information.

### pH pulsing assay

pH pulsing assay was adapted for our microscope as described previously^5^. Briefly, FR-AGS cells were transfected with appropriate constructs. The pH pulsing assay was carried out in a custom designed perfusion set up mounted on a TIRF microscope. The buffer exchanged occurred every 3s at 30°C and end of each exchange sequential images were taken. See S.I. for further details.

### Endocytic assay and microscope details

Endocytic assays were carried out as previously described^15^ with minor modifications. HRP uptake and ultrastructure analysis of endocytic structures using electron microscopy were carried out as previously described^18^. See S.I. for further details of the assay and microscopes used.

### GBP-APEX labelling

GBP-APEX labelling of GFP-IRSp53 was carried out as described previously^50^. See S.I. for further details.

### Analysis

In all cases images were analyzed with ImageJ and/or custom software written in MATLAB (The Mathworks, Natick, Massachusetts, United States). The number of cells and repeats of the experiments are mentioned in the legends and figures. Statistical significance (*p)* was calculated by Mann-Whitney U test and 2 sample student’s t-test, as reported in the legends.

### Code availability

Codes are available upon request. The algorithm is described in S.I.

### pH pulsing assay analysis

A semi-automated analysis was developed on MATLAB to identify newly formed endocytic vesicles in the pH pulsing assay and trace their intensity over time in pH 7, pH 5 and RFP channels. The traces are average of many individual traces of all the endosomes pooled from different cells, which are compared with randomly placed spots within the cells. See S.I. for further details.

**Supplementary Information** is provided as a separate file.

## Acknowledgments

We thank Ramya Purkanti, Kabir Hussain and Balaji Ramalingam for help with the analysis, Neeraj Sebastian for making TagRFPt-CDC42, R.P. for her help with BAR domain database creation, Marcus Taylor (UCSF and NCBS) for mCherry-IRSp53 and SecGFP-TfR constructs, Gero Miesenbӧck (University of Oxford) for ecliptic-GPI, Paul Melançon (University of Alberta) for ARF1-mCherry, Catherine Jackson (National Institutes of Health) for GBF1-mCherry, Roland Wedlich-Soeldner (Universität Münster) for pRuby-lifeact, TagRFPt PICK1 and GFP-PICK1 (Harvey McMohan, MRC), Mike Way (The Francis Crick Institute) for GFP-NWASPΔVCA and GFP-NWASP-VCA domain, J. Hanley (University of Bristol, UK) for pIRES-EGFP-PICK KK-EE mutant and N-WASP CA domain and Manoj Matthew from CIFF (Central Imaging and Flow cytometry Facility, NCBS) for helping to set up the pH pulsing setup. Authors were supported by Wellcome Trust-DBT India Alliance Early Career Fellowship (G.M.), NCBS-TIFR graduate student fellowship (M.S.). J.C. Bose Fellowship (S.M.), Wellcome Trust-DBT India Alliance Intermediate Fellowship and Simons Centre for Living Machines (M.T.), and grants, DBT-CoE (S.M), R.G.P was supported by the National Health and Medical Research Council (NHMRC) of Australia (program grant, APP1037320 and Senior Principal Research Fellowship, 569452), and the Australian Research Council Centre of Excellence in Convergent Bio-Nanoscience and Technology (CE140100036), G.S. and A.D. were supported by Italian Association for Cancer Research Investigator Grant (10168 and 18621 to G.S.) and European Research Council (268836 to G.S.). We acknowledge the Australian Microscopy & Microanalysis Research Facility at the Centre for Microscopy and Microanalysis at The University of Queensland. We acknowledge the CIFF (Central Imaging and Flow cytometry Facility at National Centre for Biological Sciences, TIFR, India. We thank K. Joseph Matthew (S.M. Laboratory) for making the schematic in Figure 1a and Figure 8.

## Author contributions

G.M. and M.S. executed and analysed all fluorescence microscopy experiments. M.S. and M.T developed the analysis. R.G.P. and J.R. performed, analysed and interpreted all electron microscopy experiments. A.D and G.S helped with IRSp53 knockout and mutant addback cell line construction. G.M., M.S. and S.M. planned all experiments and wrote the manuscript. All authors contributed to the writing of the manuscript.

## Author information

Correspondence and request for the materials should be addressed to S.M.

## Supplementary Figure Legends

**Figure S1:**
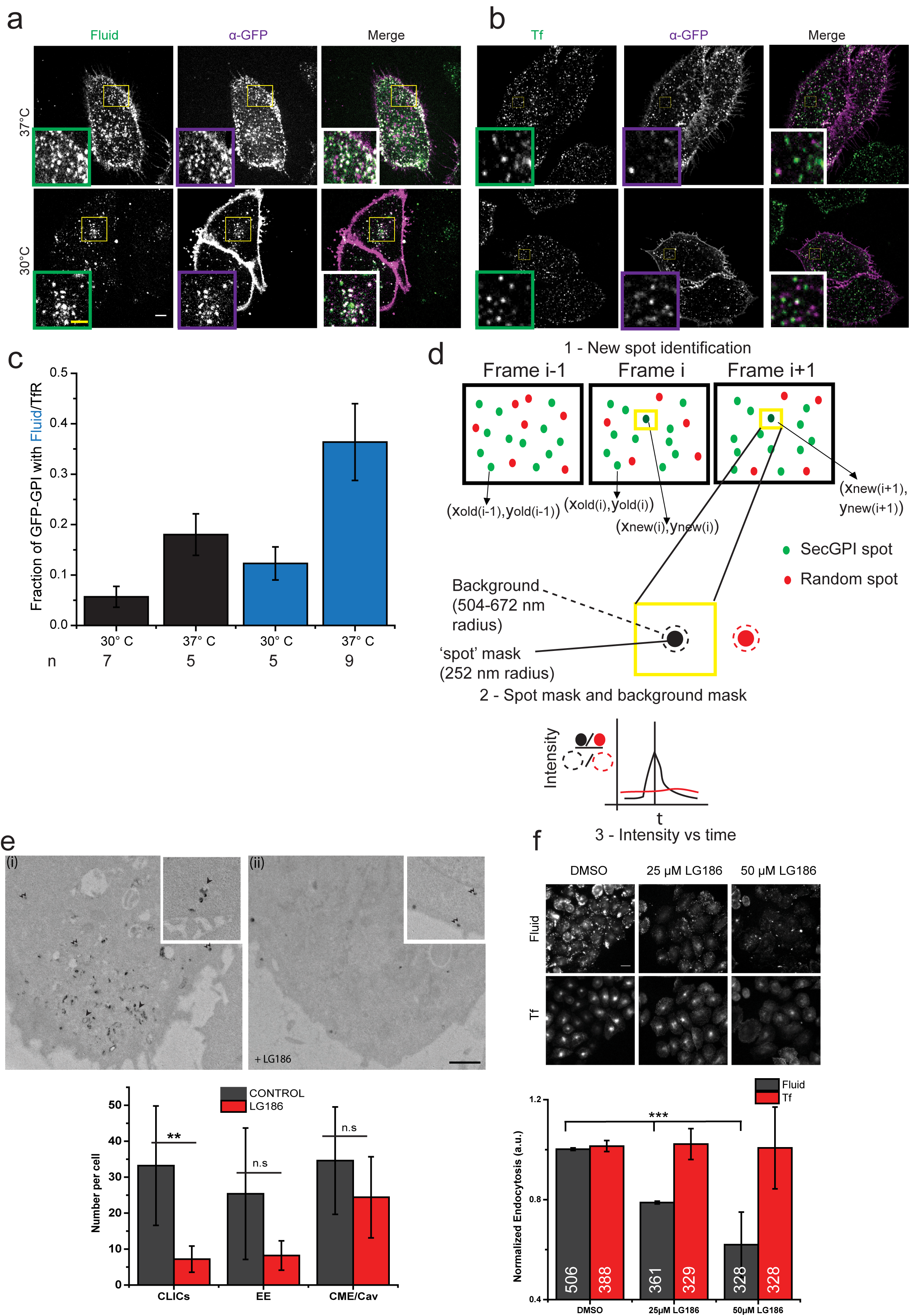
Characterization of endocytosis in AGS cells. (**a-b**) Endocytic route of SecGFP-GPI in AGS cells. Single confocal plane shows a 3 minute pulse of α-GFP Fab [**a-b**: magenta in Merge] at 37°C (upper panel) and 30°C (lower panel) along with TMR-dextran (**a**: green in Merge) or A647-Tf (**b**: green in Merge) in AGS cells transiently transfected with SecGFP-GPI. Insets show a magnified view of the marked areas. (**c**) Plot (top) showing quantification of the fraction of GFP-GPI endocytic vesicles containing fluid or Tf along. The number of cells is shown below the graph. (**d**) Schematic of pH pulsing analysis, steps (1-3) used for identifying and quantifying the fluorescence spots associated with newly formed endocytic vesicles in the sequential frames of the pH 5 montage. **Step 1** - New spot identification: Each spot (green) in i^th^ frame is compared with the previous frame and is considered new if no nearest neighbor is found by euclidean distance search within 5 pixels (1 pixel = 84nm). Random (red) are generated randomly within the cell mask. **Step 2** - Spot mask (green filled) and background mask (green dashed) considered around the centroid the new spot identified in the step - 1 [also see pH 5 frame in the middle panel of (**Figure 1a**)]. The process is repeated for Random spot mask (red filled) and background mask (red dashed). **Step 3** - schematic of the temporal profile of a new spot (black) and a random spot. Multiple traces pooled from different spots and cells over different days is averaged and shown in rest of the figures. X-axis represents time and Y-axis represents fold change in intensity in the spot over the background. See S.I. for detailed description. (**e**) Untreated AGS (Control, (i)) or LG186-treated AGS (ii) were incubated for 2 minute at 37°C with 10mg/ml HRP as a fluid phase marker before processing for electron microscopy. Endocytic structures close to the plasma membrane (PM) are filled with the electron dense peroxidase precipitate. Control cells show a range of endocytic structures including vesicular structures (CCP/Cav & EE) (a pair of small arrowheads) and tubular/ring-shaped putative CLIC/GEECs (large arrowhead) but the drug-treated cells show predominant labeling of vesicular profiles. Histogram shows mean endocytic structures quantified per cell (n = 5). CCP/Cav represent vesicles derived from clathrin mediate or caveolar endocytosis and EE represent early endosomes (See S.I.). (**f**) Histogram shows quantification of 5 minute fluid-uptake in AGS cells when treated with indicated concentrations of LG186 or DMSO. Data is pooled from 2 independent experiments and the number of cells indicated below the graph. Scale bar, 5µm (**a-b**), 4µm (**a-b, inset**), 1 µm (**e**) & 20µm (**f**) respectively. Error bars (**c**) represent s.e.m. and (**e-f**) s.d. respectively. *p-value* < 0.001 (∗∗) 2-sample student’s T-test (**e**) and MannWhitney U test (**f**).

**Figure S2:**
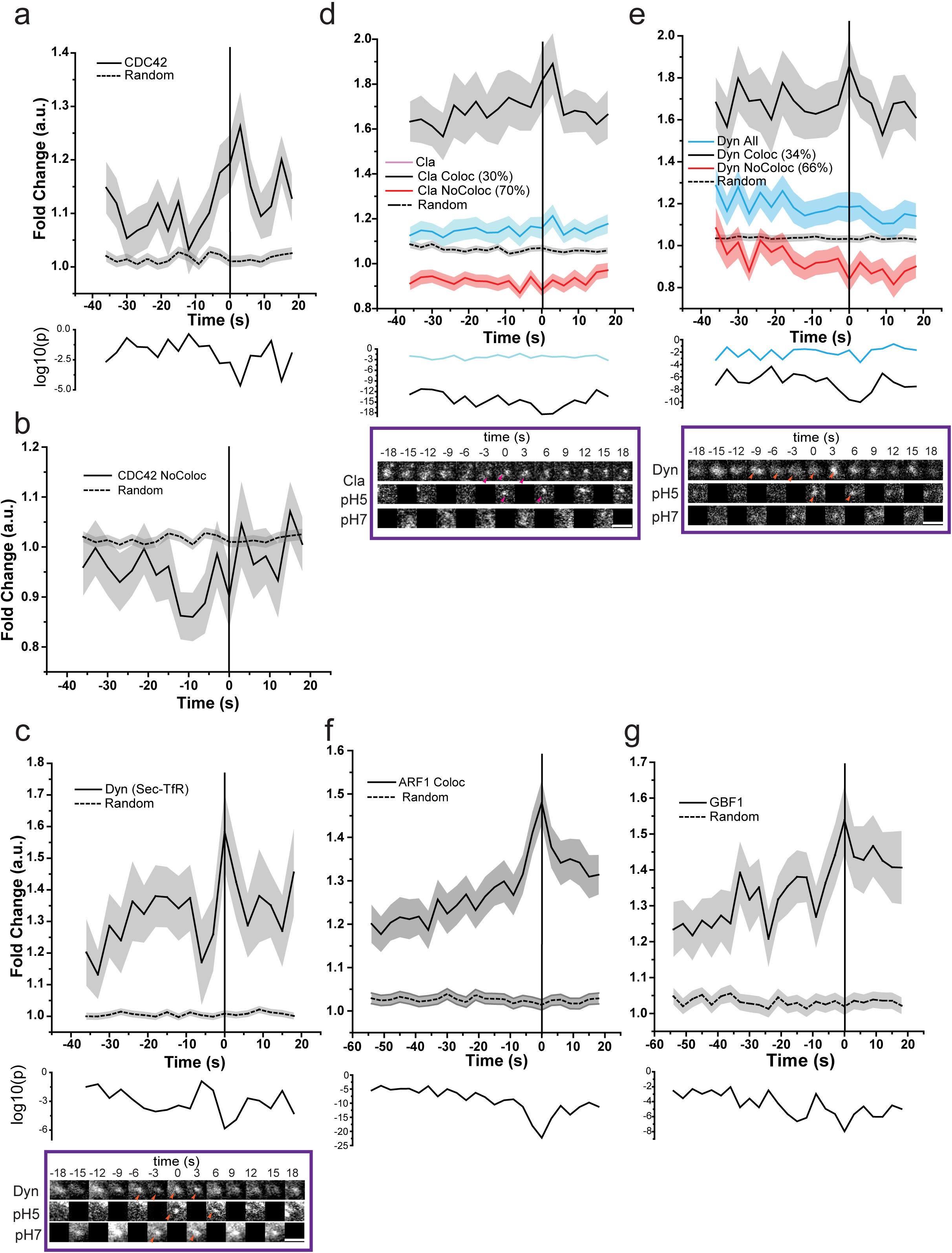
Characterization of pH pulsing assay for visualizing SecGFP-GPI endocytic vesicle formation. (**a-b**) Graphs show the average normalized fluorescence intensity versus time traces for the recruitment of TagRFPt-CDC42 (CDC42 All) to all forming SecGFP-GPI endocytic sites (**a**) or endocytic sites that do not co-detect TagRFPt-CDC42 (CDC42 NoColoc; 44%) and its corresponding random intensity trace (n, Table 2). (**c-g**) Graphs shows the average normalized fluorescence intensity versus time trace for the recruitment of mCherry-dynamin to forming SecTfR endocytic sites. [**c**; (n = 21 SecTfR and 1448 random spots from 6 cells, 2 experiments), mCherry-clathrin to forming SecGFP-GPI endocytic sites [**d**; n, Table 2], mCherry-dynamin [**e**; n, Table 2] to forming SecGFP-GPI endocytic sites, mCherry-ARF1 to forming SecGFP-GPI endocytic sites [**f**; (n = 228, as in **Figure 1e**); Note the extended time axis], mCherry-GBF1 to forming SecGFP-GPI endocytic sites [**g**; (n = 75, as in **Figure 1e**); Note the extended time axis], and their corresponding random intensity trace. A representative montage depicted below (**c-e**). Arrowheads indicate the spot. The random traces were derived from randomly assigned spots of the same radius as the endocytic regions, as detailed in S.I. Endocytic distribution at each time point was compared to the random distribution by Mann-Whitney U test and the log_10_ (p) [log_10_ (0.05) is −1.3 and log_10_ (0.001) is −2.5] is plotted below each trace (**a & c-f)**. Error bars represent s.e.m. for (**a & c-f**). Scale bar, 1.5µm (**c-e**).

**Figure S3:**
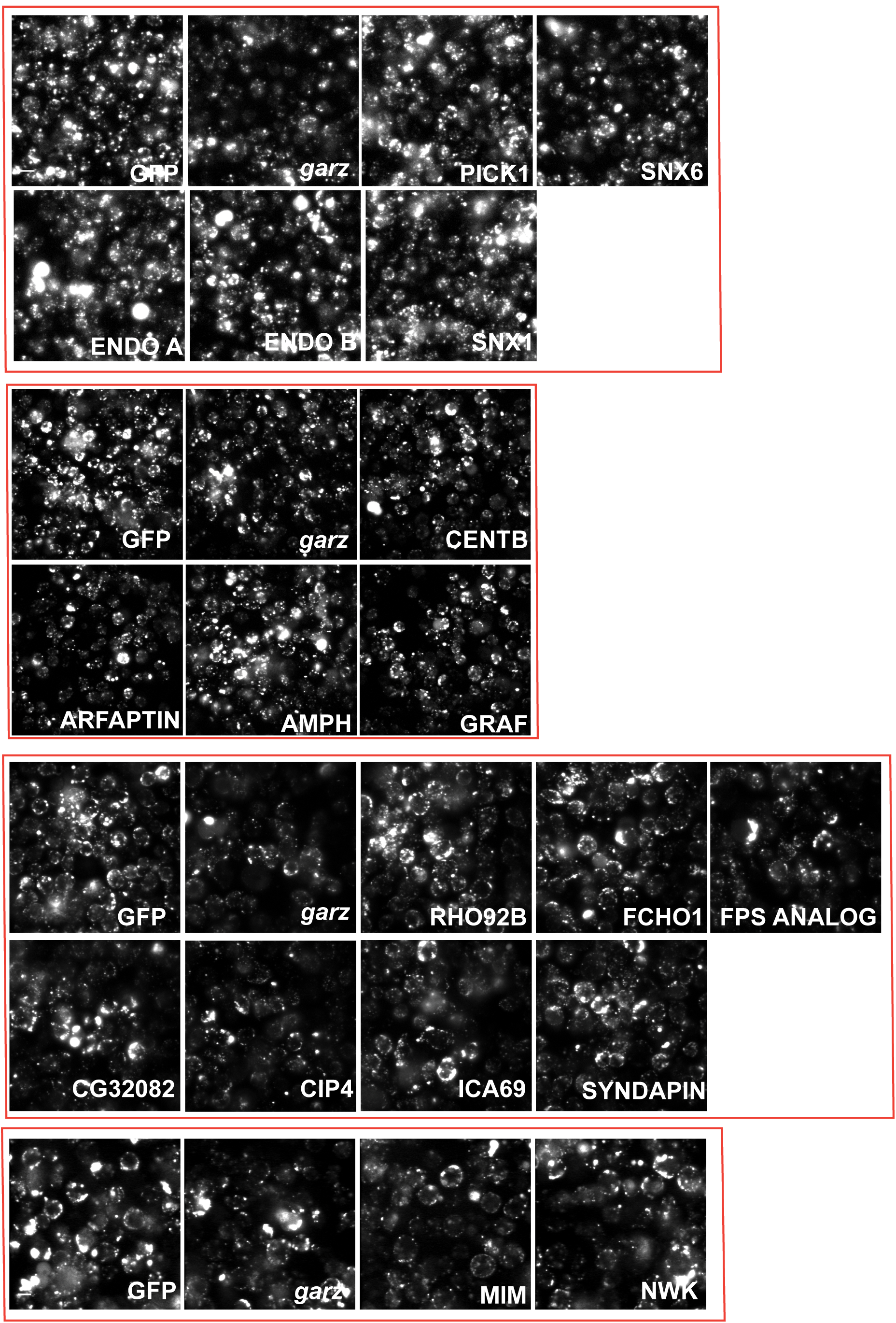
RNAi screen reveals BAR domain proteins involved in CG endocytosis. Representative images for data shown in Figure 2b. Scale bar is 20µm.

**Figure S4:**
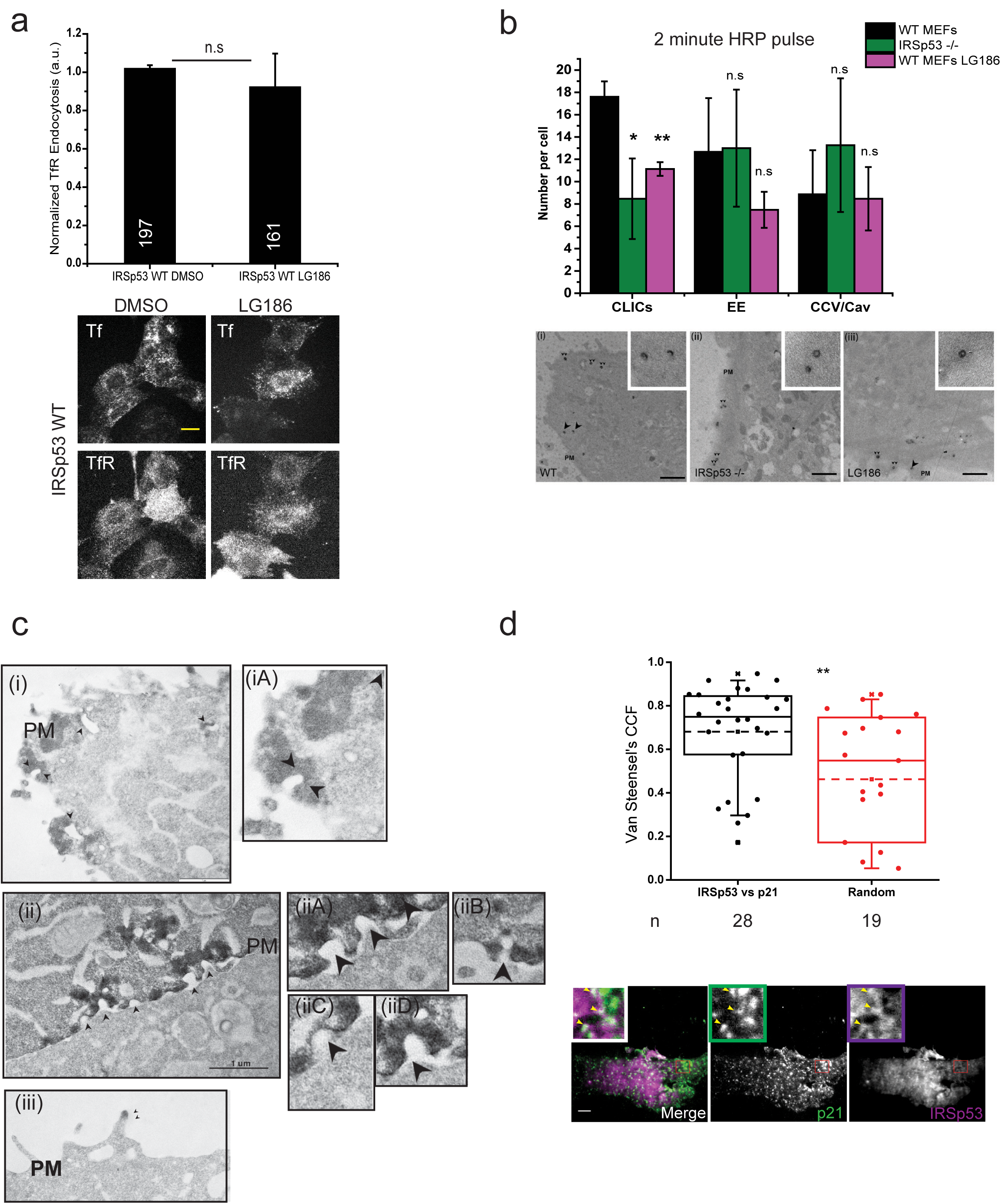
IRSp53 is involved in CG endocytosis. (**a**) Histogram (top) showing quantification of mean 5 minute normalized TfR uptake in IRSp53 WT cells when treated with 10µM LG186 for 45 minutes along with the representative images (bottom). Data is pooled from 2 independent experiments and the number of cells is indicated in the figure. (**b**) Histogram (top) shows mean number of endocytic structures per cell from the electron microscope images (below). Data pooled from 3 independent blocks with 5 cells each. Untreated WT MEFs (WT, i), IRSp53 null MEFs (IRSp53-/-, ii) or LG186-treated WT MEFs (LG186, iii) were incubated for 2 minute at 37°C with 10mg/ml HRP as a fluid phase marker before processing for electron microscopy. Endocytic structures close to the plasma membrane (PM) are filled with the electron dense peroxidase precipitate. WT cells (left) show a range of endocytic structures including vesicular structures (double arrowheads) and tubular/ring-shaped putative CLIC/GEECs (large arrowheads) but the KO cells (middle) and LG186-treated (right) cells show predominant labeling of vesicular profiles. (**c**) Electron micrographs of AGS cells co-transfected-GFP-IRSp53 and GBP-Apex. (**i-ii**) Reaction product is highly patched (arrowheads) on the plasma membrane (PM) along with zoomed regions on the right. (**iii**) Double arrowheads indicate specific labeling within defined microdomains of filopodia. (**d**) Plot (top) showing quantification of co-localizing IRSp53 with p21 (ARP2/3 complex subunit) using ImageJ plugin (Van Steensel’s CCF, See S.I.) when compared with its random. Representative (bottom) images of AGS cells co-expressing mEmerald-p21 subunit with mCherry-IRSp53 which were fixed and imaged with TIRFM, with zoomed inset at top left corner. Data is pooled from 2 independent experiments and the number of cells is indicated below the graph. Scale bar, 20µm (**a**), 1µm (**b-c**), 5µm (**d**) respectively. *p-value* < 0.01 (∗), and 0.001(∗∗) by Mann-Whitney U Test (**d**) and 2-sample student’s T-test (**b**).

**Figure S5:**
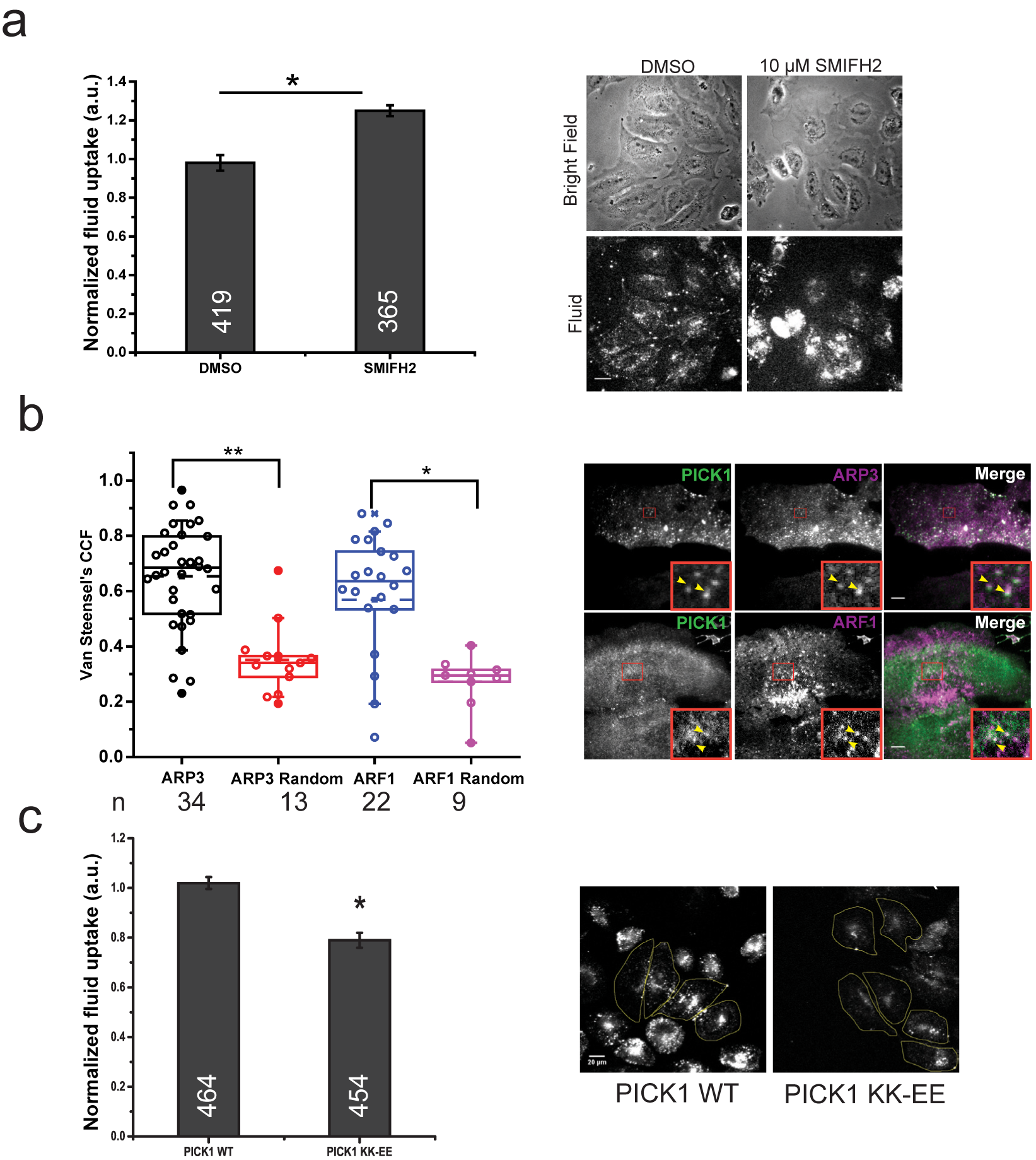
ARP2/3 is negatively regulated by PICK1 for CG endocytosis independent of N-WASP. (**a**) Histogram (left) showing quantification of 5 minute pulse fluid-phase in AGS cells treated with either DMSO or 10µM SMIFH2 along with representative images (right). Data is pooled from 2 independent experiments and the number of cells is indicated in the figure. (**b**) Plot (left) showing quantification of co-localizing PICK1 with ARP3 or ARF1 using ImageJ plugin (Van Steensel’s CCF, See S.I.) when compared with its random. Data is pooled from 2 independent experiments and the number of cells is indicated below the graph. Representative (right) images of AGS cells co-expressing GFP-ARF1 with TagRFP-PICK1 or GFP-PICK1 and mCherry-ARP3 which were fixed and imaged with TIRFM, with zoomed inset at bottom left corner. (**c**) Histogram (left) showing quantification of mean 5 minute pulse fluid-phase in AGS cells overexpressing either pIRES-PICK1 WT or PICK1 KK-EE mutant. Data is pooled from 2 independent experiments and the number of cells is indicated in the figure. Error bar represent s.d. (**a** & **c**) Scale bar, 20µm (**a** & **c**), 5µm (**b**)*. p-value* < 0.01 (∗), and 0.001 (∗∗) by Mann-Whitney U Test (**a-c**).

**Figure S6:**
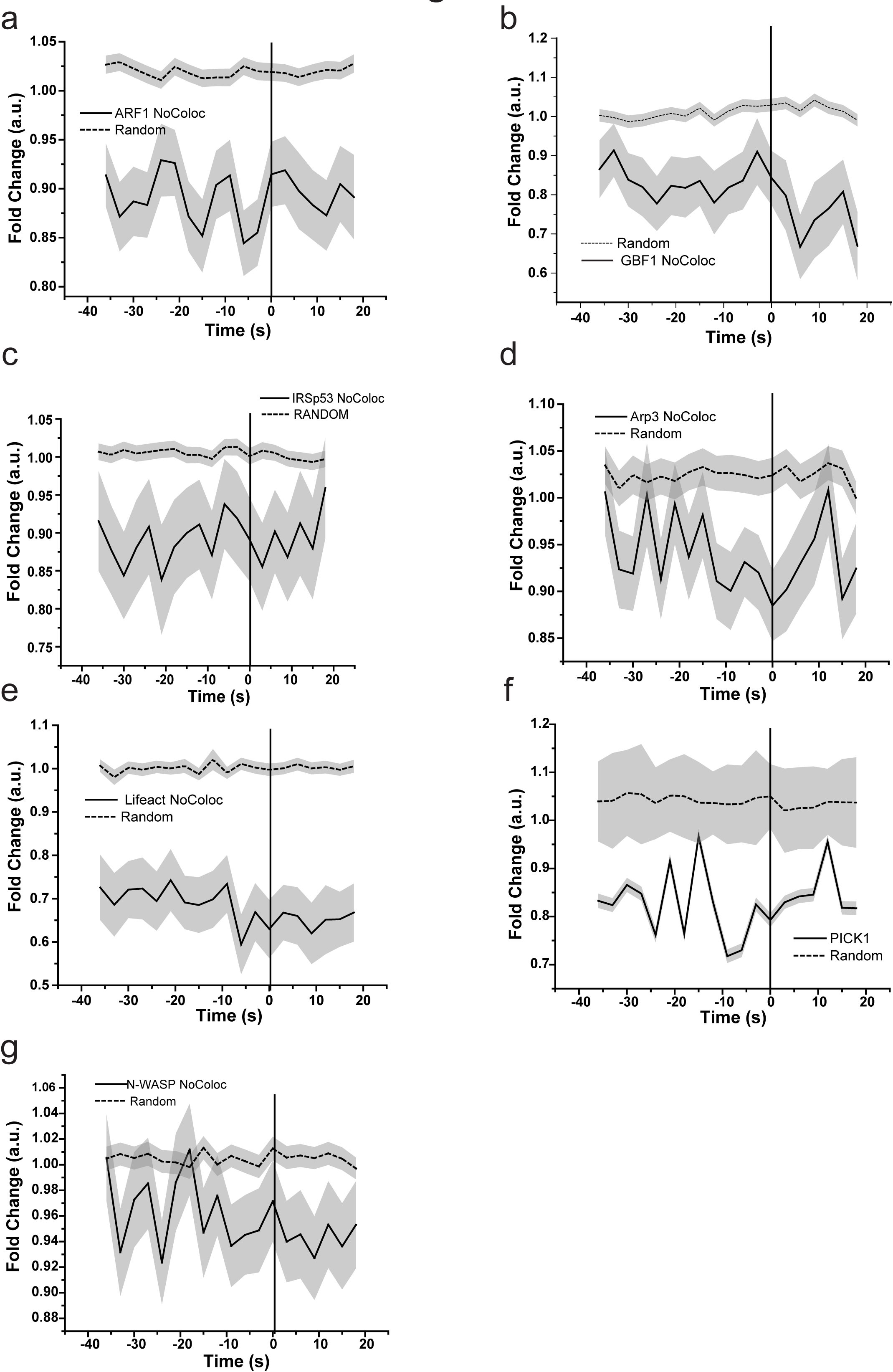
Fold change over time profiles of SecGFP-GPI spots that did not show a co-detected X-FP spot. (**a-h**) Graphs show the average normalized fluorescence intensity versus time traces for the recruitment of mCherry-ARF1 NoColoc (**a**), mCherry-GBF1 NoColoc (**b**), mCherry-IRSp53 NoColoc (**c**), mCherry-ARP3 NoColoc (**d**), pRuby-Lifeact NoColoc (**e**), TagRFPt-PICK1 NoColoc (**f**), mCherry-NWASP NoColoc (**g**) compared to their respective random trace. The random traces were derived from randomly assigned spots of the same radius as the endocytic regions, as detailed in S.I. They represent the fraction of SecGFP-GPI that failed to show a co-detected the X-FP spots. For n see Table 2 and S.I. for further information.

**Supplementary Movie 1:** Representative pH pulsing movie. AGS cells co-transfected with SecGFP-GPI and mCherry-ARF1 were imaged as described (See, S.I., pH pulsing assay). The yellow circles depict when a new event is detected. Three, movies are stitched together, where SecGFP-GPI/pH 7 (left), SecGFP-GPI/ pH 5 (middle) and mCherry-ARF1/both pH 5 & pH 7 (right) may be visualized.

**Supplementary Movie 2:** Representative event of nascent SecGFP-GPI endosome detection (zoomed from Supplementary Movie 1). AGS cells co-transfected with SecGFP-GPI and mCherry-ARF1 were imaged as described (See, S.I., pH pulsing assay). The yellow circles depict when a new event is detected. 3 movies are stitched together, SecGFP-GPI/pH 7 (left), SecGFP-GPI/ pH 5 (middle) and mCherry-ARF1/both pH 5 & pH 7 (right).

**Supplementary Movie 3:** 3D-Tomogram of APEX based detection of GFP-IRSp53 on CLICs (Representative movie 1). AGS cells co-transfected with APEX-GBP and GFP-IRSp53 were processed and imaged as described (See, S.I. Electron Microscopy). The DAB reaction was performed and the cells were processed for electron tomography (See S.I.). First half of the movie depicts sections of the original tomogram followed by density-based thresholded sections. The whole of PM of the tomographic volume was rendered and enlarged tubular regions of interest show GFP-IRSp53 recruitment.

**Supplementary Movie 4:** 3D-Tomogram of APEX based detection of GFP-IRSp53 on CLICs (Representative movie 1). AGS cells co-transfected with APEX-GBP and GFP-IRSp53 were processed and imaged as described (See, S.I. Electron Microscopy). The DAB reaction was performed and the cells were processed for electron tomography (See S.I.). First half of the movie depicts sections of the original tomogram followed by density-based thresholded sections. The whole of PM of the tomographic volume was rendered and enlarged tubular regions of interest show GFP-IRSp53 recruitment.

## REFERENCES

1. Johannes, L., Parton, R. G., Bassereau, P. & Mayor, S. Building endocytic pits without clathrin. Nat. Rev. Mol. Cell Biol. 42, 1–11 (2015).

2. Mayor, S., Parton, R. G. & Donaldson, J. G. Clathrin-independent pathways of endocytosis. Cold Spring Harb. Perspect. Biol. 6, (2014).

3. Kirchhausen, T. Imaging endocytic clathrin structures in living cells. Trends Cell Biol. 19, 596–605 (2009).

4. Kirchhausen, T., Owen, D. & Harrison, S. C. Molecular structure, function, and dynamics of clathrin-mediated membrane traffic. Cold Spring Harb. Perspect. Biol. 6, a016725–a016725 (2014).

5. Taylor, M. J., Perrais, D. & Merrifield, C. J. A high precision survey of the molecular dynamics of mammalian clathrin-mediated endocytosis. PLoS Biol. 9, e1000604 (2011).

6. Kaksonen, M., Toret, C. P. & Drubin, D. G. Harnessing actin dynamics for clathrin-mediated endocytosis. Nat. Rev. Mol. Cell Biol. 7, 404–414 (2006).

7. Cocucci, E., Gaudin, R. & Kirchhausen, T. Dynamin recruitment and membrane scission at the neck of a clathrin-coated pit. Mol Biol Cell 25, 3595–3609 (2014).

8. Shnyrova, A. V et al. Geometric catalysis of membrane fission driven by flexible dynamin rings. Science 339, 1433–6 (2013).

9. Parton, R. G. & Simons, K. The multiple faces of caveolae. Nat. Rev. Mol. Cell Biol. 8, 185–94 (2007).

10. Boucrot, E. et al. Endophilin marks and controls a clathrin-independent endocytic pathway. Nature 517, 460–5 (2015).

11. Renard, H.-F. et al. Endophilin-A2 functions in membrane scission in clathrin-independent endocytosis. Nature 517, 493–6 (2015).

12. Sabharanjak, S., Sharma, P., Parton, R. G. & Mayor, S. GPI-anchored proteins are delivered to recycling endosomes via a distinct cdc42-regulated clathrin-independent pinocytic pathway. Dev. Cell 2, 411–423 (2002).

13. Guha A, Sriram V, Krishnan KS, M. S. Shibire mutations reveal distinct dynamin-independent and -dependent endocytic pathways in primary cultures of Drosophila hemocytes. J. Cell Sci. 116, 3373–3386 (2003).

14. Chadda, R. et al. Cholesterol-sensitive Cdc42 activation regulates actin polymerization for endocytosis via the GEEC pathway. Traffic 8, 702–717 (2007).

15. Kumari, S. & Mayor, S. ARF1 is directly involved in dynamin-independent endocytosis. Nat. Cell Biol. 10, 30–41 (2008).

16. Gupta, G. D. et al. Population distribution analyses reveal a hierarchy of molecular players underlying parallel endocytic pathways. PLoS One 9, (2014).

17. Gupta, G. D. et al. Analysis of endocytic pathways in Drosophila cells reveals a conserved role for GBF1 in internalization via GEECs. PLoS One 4, (2009).

18. Howes, M. T. et al. Clathrin-independent carriers form a high capacity endocytic sorting system at the leading edge of migrating cells. J. Cell Biol. 190, 675–691 (2010).

19. Bitsikas, V. et al. Clathrin-independent pathways do not contribute significantly to endocytic flux. Elife 3, e03970 (2014).

20. Römer, W. et al. Actin dynamics drive membrane reorganization and scission in clathrin-independent endocytosis. Cell 140, 540–53 (2010).

21. Mains, P. E., Sulston, I. A. & Wood, W. B. Dominant maternal-effect mutations causing embryonic lethality in Caenorhabditis elegans. Genetics 125, 351–369 (1990).

22. Kirkham, M. et al. Ultrastructural identification of uncoated caveolin-independent early endocytic vehicles. J. Cell Biol. 168, 465–476 (2005).

23. Nonnenmacher, M. & Weber, T. Adeno-associated virus 2 infection requires endocytosis through the CLIC/GEEC pathway. Cell Host Microbe 10, 563–576 (2011).

24. Kalia, M. et al. Arf6-independent GPI-anchored protein-enriched early endosomal compartments fuse with sorting endosomes via a Rab5/phosphatidylinositol-3’-kinase-dependent machinery. Mol. Biol. Cell 17, 3689–704 (2006).

25. Kettle, E. et al. A Cholesterol-Dependent Endocytic Mechanism Generates Midbody Tubules During Cytokinesis. Traffic 16, 1174–1192 (2015).

26. Miesenböck, G., De Angelis, D. A. & Rothman, J. E. Visualizing secretion and synaptic transmission with pH-sensitive green fluorescent proteins. Nature 394, 192–195 (1998).

27. Merrifield, C. J., Perrais, D. & Zenisek, D. Coupling between clathrin-coated-pit invagination, cortactin recruitment, and membrane scission observed in live cells. Cell 121, 593–606 (2005).

28. Merrifield, C. J., Feldman, M. E., Wan, L. & Almers, W. Imaging actin and dynamin recruitment during invagination of single clathrin-coated pits. Nat Cell Biol 4, 691–698 (2002).

29. Qualmann, B., Koch, D. & Kessels, M. M. Focus Review Let’s go bananas: revisiting the endocytic BAR code Cellular membranes—between relaxation and defined topology Shaping membranes with BAR domains. EMBO J. 30, 3501–3515 (2011).

30. Frost, A., Unger, V. M. & De Camilli, P. The BAR Domain Superfamily: Membrane-Molding Macromolecules. Cell 137, 191–196 (2009).

31. Lundmark, R. et al. The GTPase-Activating Protein GRAF1 Regulates the CLIC/GEEC Endocytic Pathway. Curr. Biol. 18, 1802–1808 (2008).

32. Kumari, S. Dynamin-independent endocytosis : molecular mechanisms and membrane dynamics Dynamin-independent endocytosis : molecular mechanisms and membrane dynamics. Tata Institute of Fundamental Research (2008).

33. Gillingham, A. K. & Munro, S. The small G proteins of the Arf family and their regulators. Annu. Rev. Cell Dev. Biol. 23, 579–611 (2007).

34. Boal, F. et al. LG186: An Inhibitor of GBF1 Function that Causes Golgi Disassembly in Human and Canine Cells. Traffic 11, 1537–1551 (2010).

35. Wassmer, T. et al. A loss-of-function screen reveals SNX5 and SNX6 as potential components of the mammalian retromer. J. Cell Sci. 120, 45–54 (2007).

36. Dai, J. et al. ACAP1 promotes endocytic recycling by recognizing recycling sorting signals. Dev. Cell 7, 771–776 (2004).

37. Hartig, S. M. et al. The F-BAR protein CIP4 promotes GLUT4 endocytosis through bidirectional interactions with N-WASp and Dynamin-2. J. Cell Sci. 122, 2283–2291 (2009).

38. Rodal, A. A., Motola-Barnes, R. N. & Littleton, J. T. Nervous Wreck and Cdc42 Cooperate to Regulate Endocytic Actin Assembly during Synaptic Growth. J. Neurosci. 28, 8316–8325 (2008).

39. Scita, G., Confalonieri, S., Lappalainen, P. & Suetsugu, S. IRSp53: crossing the road of membrane and actin dynamics in the formation of membrane protrusions. Trends Cell Biol. 18, 52–60 (2008).

40. Nakagawa, H. et al. IRSp53 is colocalised with WAVE2 at the tips of protruding lamellipodia and filopodia independently of Mena. J. Cell Sci. 116, 2577–83 (2003).

41. Goh, W. I. et al. mDia1 and WAVE2 proteins interact directly with IRSp53 in filopodia and are involved in filopodium formation. J. Biol. Chem. 287, 4702–4714 (2012).

42. Miki, H., Yamaguchi, H., Suetsugu, S. & Takenawa, T. IRSp53 is an essential intermediate between Rac and WAVE in the regulation of membrane ruffling. Nature 408, 732–735 (2000).

43. Disanza, A. et al. CDC42 switches IRSp53 from inhibition of actin growth to elongation by clustering of VASP. EMBO J. 32, 2735–50 (2013).

44. Vaggi, F. et al. The Eps8/IRSp53/VASP network differentially controls actin capping and bundling in filopodia formation. PLoS Comput. Biol. 7, (2011).

45. Disanza, A. et al. Regulation of cell shape by Cdc42 is mediated by the synergic actin-bundling activity of the Eps8–IRSp53 complex. Nat. Cell Biol. 8, 1337–1347 (2006).

46. Funato, Y. et al. IRSp53/Eps8 complex is important for positive regulation of Rac and cancer cell motility/invasiveness. Cancer Res. 64, 5237–44 (2004).

47. Kast, D. J. et al. Mechanism of IRSp53 inhibition and combinatorial activation by Cdc42 and downstream effectors. Nat. Struct. Mol. Biol. 21, 413–22 (2014).

48. Lewis-Saravalli, S., Campbell, S. & Claing, A. ARF1 controls Rac1 signaling to regulate migration of MDA-MB-231 invasive breast cancer cells. Cell. Signal. 25, 1813–1819 (2013).

49. Prévost, C. et al. IRSp53 senses negative membrane curvature and phase separates along membrane tubules. Nat. Commun. 6, 8529 (2015).

50. Ariotti, N. et al. Modular Detection of GFP-Labeled Proteins for Rapid Screening by Electron Microscopy in Cells and Organisms. Dev. Cell 35, 513–525 (2015).

51. Kim, B. L. et al. The Cdc42 effector IRSp53 generates filopodia by coupling membrane protrusion with actin dynamics. J. Biol. Chem. 283, 20454–20472 (2008).

52. Havrylenko, S. et al. WAVE binds Ena/VASP for enhanced Arp2/3 complex–based actin assembly. Mol. Biol. Cell 26, 55–65 (2015).

53. Nolen, B. J. et al. Characterization of two classes of small molecule inhibitors of Arp2/3 complex. Nature 460, 1031–4 (2009).

54. Rizvi, S. A. et al. Identification and characterization of a small molecule inhibitor of formin-mediated actin assembly. Chem. Biol. 16, 1158–68 (2009).

55. Riedl, J. et al. Lifeact: a versatile marker to visualize F-actin. Nat. Methods 5, 605–7 (2008).

56. Cerione, R. A. Cdc42: New roads to travel. Trends Cell Biol. 14, 127–132 (2004).

57. Rohatgi, R. et al. The interaction between N-WASP and the Arp2/3 complex links Cdc42-dependent signals to actin assembly. Cell 97, 221–231 (1999).

58. Rocca, D. et al. The Small GTPase Arf1 Modulates Arp2/3-mediated actin polymerization via PICK1 to regulate synaptic plasticity. Neuron 79, 293–307 (2013).

59. Rocca, D. L., Martin, S., Jenkins, E. L. & Hanley, J. G. Inhibition of Arp2/3-mediated actin polymerization by PICK1 regulates neuronal morphology and AMPA receptor endocytosis. Nat. Cell Biol. 10, 259–71 (2008).

60. Nakamura, Y. et al. PICK1 inhibition of the Arp2/3 complex controls dendritic spine size and synaptic plasticity. EMBO J. 30, 719–730 (2011).

61. Thorsen, T. S. et al. Identification of a small-molecule inhibitor of the PICK1 PDZ domain that inhibits hippocampal LTP and LTD. Proc. Natl. Acad. Sci. 107, 413–418 (2010).

62. Sandvig, K., Torgersen, M. L., Raa, H. A. & Van Deurs, B. Clathrin-independent endocytosis: From nonexisting to an extreme degree of complexity. Histochem. Cell Biol. 129, 267–276 (2008).

63. Jolla, L. et al. Clathrin-independent Pinocytosis Is Induced in Cells Overexpressing a Temperature-sensitive Mutant of Dynamin. J. Cell Biol. 131, 69–80 (1995).

64. Kirkham, M. & Parton, R. G. Clathrin-independent endocytosis: New insights into caveolae and non-caveolar lipid raft carriers. Biochim. Biophys. Acta - Mol. Cell Res. 1745, 273–286 (2005).

65. Kaksonen, M., Toret, C. P. & Drubin, D. G. A modular design for the clathrin- and actin-mediated endocytosis machinery. Cell 123, 305–320 (2005).

66. Takeya, R., Takeshige, K. & Sumimoto, H. Interaction of the PDZ domain of human PICK1 with class I ADP-ribosylation factors. Biochem. Biophys. Res. Commun. 267, 149–155 (2000).

67. Simunovic, M. et al. Friction Mediates Scission of Tubular Membranes Scaffolded by BAR Proteins. Cell 170, 172–184.e11 (2017).

